# Protein-specific crowding accelerates aging in phase-separated droplets

**DOI:** 10.1101/2023.12.10.570970

**Authors:** Mateusz Brzezinski, Pablo G. Argudo, Tom Scheidt, Miao Yu, Edward A. Lemke, Jasper J. Michels, Sapun H. Parekh

## Abstract

Crowding agents, such as polyethylene glycol (PEG, are often used to mimic the cellular cytoplasm in protein assembly studies. Despite the perception that crowding agents have an inert nature, recent work has shown they are not bystanders while proteins interact. Here, we explore the diverse effects of PEG on the phase separation and maturation of proteins. We use two proteins, the FG domain of Nup98 and bovine serum albumin (BSA), which represent an intrinsically disordered protein and a protein with well-established secondary structure, respectively. PEG expedites the maturation of Nup98, enhancing denser protein packing and fortifying hydrophobic interactions which hasten beta-sheet formation and subsequent droplet gelation. In contrast for BSA, PEG appears to enhance droplet stability and limits available solvent for the protein solubilization, without inducing significant changes to the secondary structure, pointing towards a significantly different behavior of the crowding agent. Interestingly, we detect almost no presence of PEG in Nup droplets whereas PEG is detectable within BSA droplets. Our findings demonstrate a nuanced interplay between crowding agents and proteins. PEG can accelerate protein maturation in LLPS systems but its partitioning and effect on protein structure in droplets is protein specific. This suggests that crowding phenomena are specific to each protein-crowding agent pair.

## INTRODUCTION

The study of proteins and their interactions is a cornerstone of modern biochemistry and molecular biology. One emerging phenomenon in this field is phase separation (PS), which appears to underlie, at least in part, the formation of membrane-less organelles (MLOs) inside the cells [1], [2]. This process in protein base systems can be driven by multivalent protein-protein interactions, leading to the formation of droplet-like structures that compartmentalize biomolecules without the need for a surrounding lipid barrier. Crucially, these protein interactions and the ensuing PS can be heavily influenced by the crowded nature of the cellular environment [3]. In this context, the role of crowding agents, which mimic the dense intracellular milieu, is being increasingly recognized and investigated [4]. Crowding agents affect PS through several key mechanisms [5]. One effect is volume exclusion, a fundamental principle underpinning their function. Volume exclusion promotes PS by reducing the available volume for other macromolecules, thereby inducing segregation and formation of distinct phases [6], [7]. Additionally, crowding agents decrease the diffusion path of proteins, enhancing opportunities for interactions and PS [8]. Lastly, by inducing a dehydration effect, crowding agents strip away the water molecules that may otherwise be associated with biomolecules, making protein-protein interactions more favorable and further promoting PS [9]. Molecular partitioning of crowding agents follows one of two pathways: competitive (segregative) and cooperative (associative) [10]–[12]. In segregative PS, protein and crowding agent macromolecules partition into distinct phases to minimize interactions. In contrast, in associative PS, both protein and crowding agent macromolecules coalesce into a single phase, driven by mutual affinity.

Polyethylene glycol (PEG) is a widely used crowding agent in biochemical studies to mimic the intracellular environment. It is a variable molecular weight polymer composed of repeating ethylene oxide units. PEG was considered an “inert” crowding agent because it affects the biomacromolecules in its vicinity primarily through physical, not chemical, interactions [13], [14]. PEG is hydrophilic and, thus, incapable of forming specific or strong interactions with proteins or other biomolecules [15]–[18]. Its presence predominantly alters the physical environment. Owing to these properties, PEG has been used to effectively mimic the crowded nature of a typical cell interior without introducing unnecessary complications from specific non-covalent interactions [4]. Recent research indicates a shift in our understanding of PEG’s role in PS. Contrary to its traditional classification as an ‘inert’ crowding agent, newer studies suggest that PEG can co-localize within protein condensates and actively participate in an associative PS [12], [19]. This highlights PEG’s potential contribution to the formation and characteristics of the resulting phase-separated structures.

Nucleoporins (Nups), key components of the nuclear pore complex (NPC), belong to the family of intrinsically disordered proteins (IDPs) that lack a well-defined structure in their native state, providing them with exceptional conformational and structural flexibility [20]. Within the highly crowded environment of the NPC, these Nups play an indispensable role in controlling molecular traffic between the nucleus and the cytoplasm [21]. Under certain conditions, Nups can undergo further transitions, leading to fibrillation, where they self-assemble into fiber-like structures *in vitro* [22], [23]. This transition is thought to be propelled by hydrophobic interactions via phenylalanine/glycine (FG) repeat motifs, suggesting a key role for non-polar forces in driving the transition towards fibrillation. The potential sequence of transitions—liquid to gel to fibrils—showcases the adaptability of Nups and their intricate contribution to cellular function [24]. However, this uncontrolled aggregation and subsequent deposition of other IDP families such as FUS (FUsed in Sarcoma) or TAU (Tubulin Associated Unit) can disrupt normal cellular functions and is associated with a range of neurodegenerative disorders, including Alzheimer’s, Parkinson’s, and Huntington’s diseases [25]–[29]. The self-assembly of these proteins is often driven by beta-sheet formation, which allows stable aggregates to form via an increase in intermolecular hydrogen bonding facilitated by the exposure of hydrophobic regions [30], [31]. Consequently, understanding the influence of the nucleoporin environment on such a structural transition of Nups is of paramount importance. This knowledge will provide critical insights into the mechanistic pathways of protein aggregation and its role in disease [32]. Numerous studies have explored the impact of PEG and other crowding agents on many proteins, spanning both IDPs and those with well-defined secondary structures [33], [34]. However, the influence of crowding on the kinetics of the structural maturation in PS-formed protein condensates remains incomplete and warrants further investigation [35], [36].

In the present investigation, we delve into the intricacies of the kinetic aspect of protein condensate maturation, focusing specifically on the structural transformation of initially liquid-like protein condensates in the presence and absence of a PEG crowding agent. We implement broadband coherent anti-Stokes Raman scattering (BCARS) and Fluorescence Recovery After Photobleaching (FRAP) to meticulously examine the ‘aging’ states of Nup98 under varying crowding conditions and time frames. Additionally, we include a comparison with PS of a solution of Bovine Serum Albumin (BSA), a well-studied protein with a defined folding. Through this study, we aim to shed light on the specifics of how molecular crowding influences protein condensate formation and transformation or maturation, and whether this influence translates universally across both disordered and folded protein assemblies or remains specific to a certain class of proteins.

## MATERIALS AND METHODS

### Nup purification, solutions, and labeling

The FG domain of *Homo sapiens* Nup98 (spanning 1-505 amino acids, excluding the Gle2-binding domain (GLEBS; 157–213 amino acids), a structured domain interspersed between the two FG domains) was expressed in *Escherichia coli* BL21 AI cells. Detailed protocols for the expression and purification are available in the Methods section of our previous publication [37]. The purified Nup98 was concentrated to a final concentration of 165 µM using 3-kDa MWCO centrifugal filters (Merck Millipore) in a solution of 2 M GdmCl, 0.2 mM TCEP and 50 mM Tris-HCl, pH 8, with the concentration measured by Pierce™ BCA Protein Assay Kit (23227, Thermo Fisher). The proteins were flash-frozen and stored at −80 °C.

For labeling, the purified Nup98 FG domain, with single cysteine mutation (A221C), was exchanged into 4 M GdmCl, 1× PBS, and 0.1 mM EDTA, and 0.2 mM TCEP, pH 7. Labeling with Alexa Fluor™ 488 maleimide (A10254, Thermo Fisher) was performed at a molar ratio of 1:2 (dye:protein) overnight at 4°C. The reaction was quenched with 10 mM DTT in 4 M GdmCl and 1× PBS, pH 7. Unreacted dye was washed off using a 3-kDa MWCO centrifugal filter, and the labeled protein was further purified with Superdex 200. Pure fractions were selected, pooled, and concentrated, and the final concentration was measured using a Duetta absorbance spectrometer (Horiba). Protein was flash-frozen and stored at −80 °C. For all fluorescence-based investigations, the Nup98 solution was doped with 1% of a fluorescently labeled version of Nup98.

### BSA solutions and labeling

Bovine Serum Albumin (BSA) Fraction V (1126GR050, neoFroxx) was dissolved in 10xPBS buffer overnight at a constant temperature of 25°C using a Thermoblock to ensure complete dissolution. The solution was then transferred to a syringe fitted with a 0.22 µm PVDF syringe filter and subsequently filtered to remove any potential contaminants. The purified aliquots were flash-frozen and stored at -80°C until further use. The final concentration of the solution was confirmed to be 125 µM using the Pierce™ BCA Protein Assay Kit (23227, Thermo Fisher).

For fluorescent BSA, fluorescein-labeled BSA (BSA-FITC), Albumin–fluorescein isothiocyanate conjugate (A9771, Merck) was used. The lyophilized conjugate was reconstituted in 10xPBS to a concentration of 500 µM, following the procedure previously outlined. After preparation, the protein solution was flash-frozen and stored at -80°C until required. For all fluorescence-driven experiments, the standard BSA solution was enriched with 1% of the fluorescently labeled BSA.

### PEG solutions

Polyethylene Glycol (PEG) with a molecular weight of 4000 g/mol (95904, Merck) was dissolved in either 1x Transport Buffer (TB; 20 mM HEPES, 110 mM KOAc, 5 mM NaOAc, 2 mM Mg(OAc)_2_, 1 mM EGTA and 2 mM DTT, pH 7.3,) [38] (Nup98 studies) or 1xPBS (BSA studies). This was achieved through overnight incubation at 40°C with continuous stirring on a hot plate, resulting in a final concentration of 40% (w/v). This stock solution was then systematically diluted to generate a series of working concentrations suitable for our studies: 0%, 2.5%, 5%, 10%, and 30% for investigations on Nup98, and 10%, 20%, and 30% for BSA studies. All solutions were stored at -80°C.

### Condensate sample preparation and preservation

For our time-dependent experiments for each sample, a 24×60 mm, #1, non-modified glass coverslip was prepared with two parallel strips of double-sided tape, leaving an approximate gap of 15 mm. Subsequently, 2 µl of the protein solution was carefully pipetted into the gap, followed by the addition of 20 µl of the buffer with PEG. After allowing about a minute for the sample to stabilize, a 20×20 mm, #1 glass coverslip was positioned over the sample to create a seal. The top coverslip was then firmly pressed onto the double-sided tape, ensuring secure adhesion. To further enhance the seal and maintain an airtight environment, each of the four edges of the top coverslip was sealed using a quick-drying, cyanoacrylate-based adhesive (UHU, super glue). This comprehensive preparation ensured an optimal environment for our subsequent time-dependent microscopy studies, minimizing any potential evaporation or contamination. Impressively, this robust setup displayed no leakage even after a month of storage, thereby reinforcing the stability and reliability of our experimental conditions.

### Fluorescence Recovery After Photobleaching

FRAP experiments were conducted using a Leica SP8 confocal microscope with a 63×, 1.20 NA water immersion objective. We utilized a 488 nm laser line for studies involving Nup98, and both 488 nm and 496 nm laser lines for BSA experiments. All image stacks acquired maintained a consistent resolution of 512×512 pixels. The scanner was configured to a scanning frequency of 700 Hz, allowing for a cycle time of 0.743 s. A total of 165 frames were recorded for each sample: the first 5 frames were captured as “pre-bleach” images using a laser power of 5%, followed by 10 bleach frames with the laser power locally intensified to 100%, and finally 150 post-bleach frames with a laser power of 5%. Throughout the imaging process, the pinhole size was held constant at 100 nm to ensure consistency in the collected data. Zoom was also maintained within a range of 15-30x. This imaging setup allowed high-resolution and consistent data collection across all FRAP experiments. After data acquisition, a custom Python script was employed to process the collected data. This script was designed to facilitate the simultaneous analysis of four bleached Regions of Interest (ROIs) while accounting for an additional background ROI (consisting of an empty plane) and a reference ROI (comprising the entire sample plane). Following this preliminary processing, data were exported to easyFRAP, an online FRAP analysis tool [39]–[41].

### Broadband coherent anti-Stokes Raman scattering (BCARS) microscopy

Raman measurements were performed using a home-built broadband coherent anti-Stokes Raman scattering (BCARS) microscope, the detailed configuration of which has been documented in prior work [42]. The pump/probe and Stokes pulses are created in a dual-output, sub-nanosecond laser source (CARS-SM-30, Leukos). These pulses are then synchronously overlapped in space and time at the sample plane of an inverted microscope (Eclipse Ti-U, Nikon) and tightly focused onto the sample using a 0.85 NA air objective (LCPlan N, Olympus). The BCARS signal, once isolated from the excitation pulses, is focused onto the slit of a spectrograph (Shamrock 303i, Andor). This disperses the spectral components onto a cooled CCD camera (Newport DU920P-BR-DD, Andor). Samples are positioned with the coverslip facing the collector and scanned using a piezo stage (Nano-PDQ 375 HS, Mad City Labs) controlled by LabView 2015 (National Instruments) software. Following data collection, the amassed hyperspectral data undergoes processing in IgorPro (WaveMetrics). The Raman-like spectra are retrieved through a modified Kramers-Kronig transform [43]. All spectra presented in this paper were phase-retrieved using buffer alone, with no macromolecules as a reference spectrum, and any background phase is removed using a Savitzky-Golay filter with a 2^nd^ order polynomial and a window size of approximately 400 cm^-1^. This protocol allows us to acquire and interpret high-quality spectral data from our samples.

For the deconvolution of the Nup98 Amide I band, we utilized a custom Python script. This script facilitated the generation of an initial seed for the parameters of the Lorentzian peaks. We derived these initial estimates in accordance with our prior published work [44]. Upon establishing these initial parameters, we input them into the peak analyzer function of OriginLab Pro. This software further refined the deconvolution and produced the final resolution of the peaks.

To analyze BSA/PEG concentrations via CH stretch peak fitting, we employed a custom Python script based on the BCARS spectra of 4 mM BSA and 75 mM PEG reference samples for the CH region (2800 - 3100 cm^-1^). These samples were processed and measured following the same protocols as described above. Spectral fitting of phase-retrieved spectra was conducted in the range of 2820 to 3050 cm^-1^, with each spectrum normalized to the highest peak of the sample. BSA and PEG concentrations were determined by fitting the normalized spectrum to a weighted sum of the pure BSA and PEG spectra, with each component being weighted by a coefficient to best fit the experimental spectrum. The fitting coefficients were then transformed into concentrations using a reverse normalization procedure to correlate with BSA or PEG reference concentrations. To account for variations linked to different alignments and to make spectra from day-to-day quantitatively comparable, we collected Raman-like spectra of water prior to each sample measurement. This was used for subsequent rescaling, ensuring that potential deviations were minimized, and thus enabling accurate and reliable analysis.

## RESULTS AND DISCUSSION

### PEG crowding drives segregative Nup98 condensation, accelerates structural maturation, and reduces molecular mobility

Liquid-liquid phase separation of Nup98, whether in the presence or absence of PEG, results in the formation of droplet-like condensates. The size of these condensates varies with the concentration of the PEG crowding agent and the condensate population namely their location, size, and shape are non-uniform on the sample, suggesting a kinetic process at work during condensate formation (**Fig. S1**). To obtain a comparable understanding of the chemical composition of both the continuous and droplet phases across all PEG concentrations, we targeted regions in the samples where both phases (droplets and a continuous phase) were present (**Fig. 1A**) (shown for 30% (w/v) PEG). Immediately obvious from the bright field imaging are overlapping droplet structures, implying the ability to form unified condensates through droplet coalescence. This indicates the liquidity of such droplets, at least during their formation, as the structure of condensates was found to be stable later during the measurement 24 hours post droplet formation. BCARS imaging allowed us to acquire intensity maps spanning the spectral range from 700 to 3700 cm^-1^ and to correlate the spatial distribution of each phase with their respective CARS spectra to understand chemical distribution across sample plane (**Fig. 1B**). For an in-depth analysis of the diversity of chemical composition inside and outside of the condensates, we made regions of interest using a representative number of pixels for both phases based on the visual representation of the intensity map. Interestingly, these regions exhibited distinct chemical variations (**Fig. 1C**). The droplet phase exhibited a typical protein fingerprint region profile with pronounced peaks at ∼ 1670 cm^-1^ for Amide I and ∼ 1000 cm^-1^, indicative of phenylalanine residues. Additionally, the C-H stretching profile for both CH_2_ and CH_3_ groups occurred in the 2920-3030 cm^-1^ region, along with a notable peak at around 3065 cm^-1^, which comes from aromatic groups of amino acids [45]–[49]. In stark contrast, the continuous phase lacked these peaks but showed vibrations associated with PEG, if present in the mixture, or nothing when the sample was protein and buffer alone. The continuous phase with PEG and Nup98 in the sample is characterized by a triple peak in the C-H stretching vibration region of 2850-3000 cm^-1^ as well as numerous peaks in the fingerprint region correlated to C-C and C-O starching and C-C-O bending and other chain bending modes [50]–[53].

**Figure 1.**
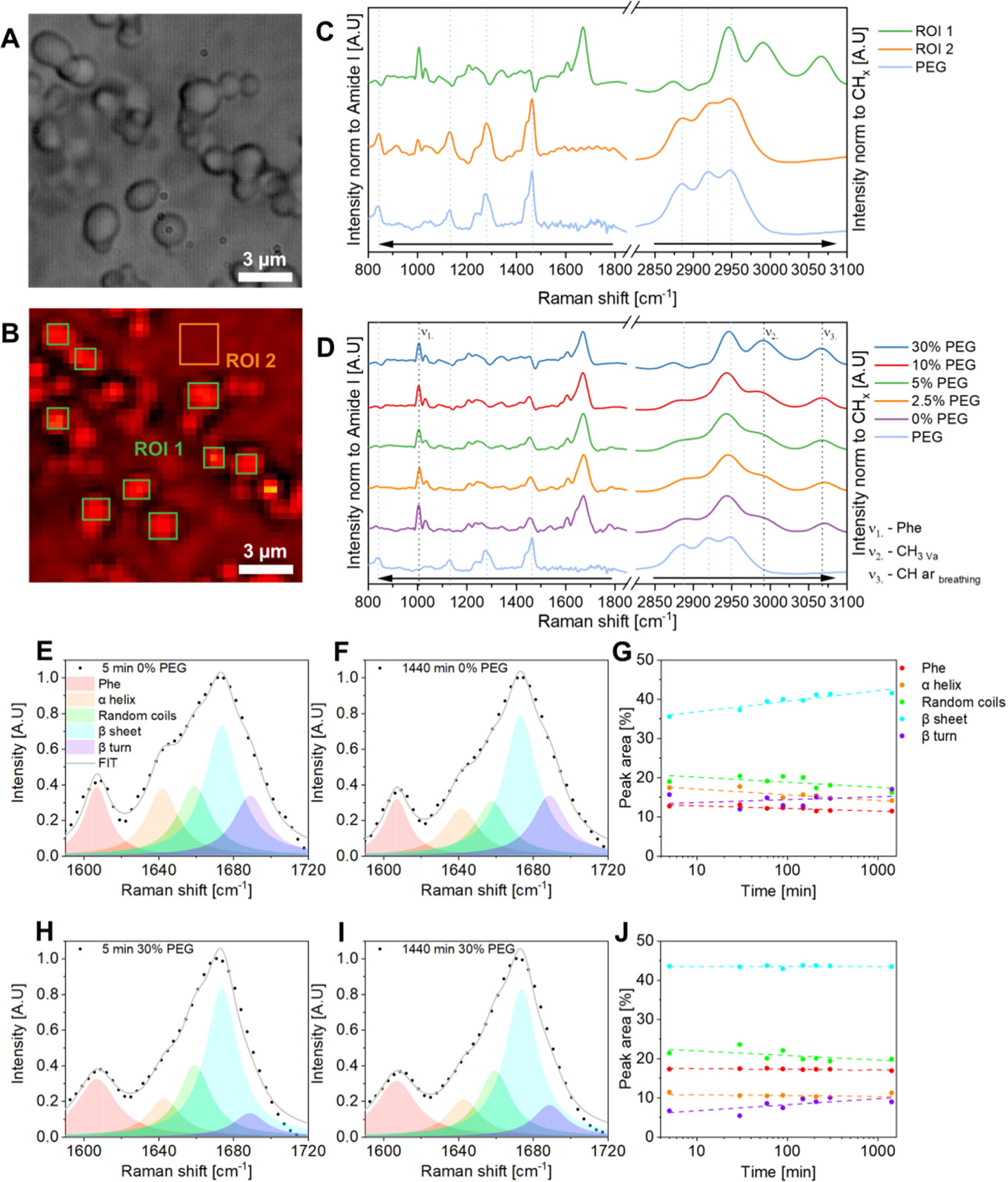
Molecular microscopy of Nup98:PEG condensate aging. In situ **A)** BF image and **B)** BCARS map imaging of the same Nup condensates formed by mixing 1:10 Nup 0.165mM with PEG 4kDa 75mM (30% m/v). Both images were captured 24h after initiation of LLPS. BCARS map shows averaged CH stretch intensity and is integrated over 2820 - 3020cm^-1^ with a pixel size of 0.375×0.375µm. On the BCARS map, ROI 1 marked green, represents a selected part of the droplet phase, and ROI 2 marked orange, represents a selected part of the continuous phase of a sample**. C)** Normalized fingerprint and CH stretch spectra comparison of ROI 1 and ROI 2 to 75mM PEG reference sample. **D)** Normalized fingerprint and CH stretch spectra of droplet phase for different PEG concentration systems measured 5min after initializing LLPS and 75mM PEG reference sample. Amide I spectra deconvolution for **E)** 0% PEG system at 5min and **F)** 24h and **H)** 30% PEG system at 5min and **I)** 24h. Thick lines are fit curves which are the sum of all subpeaks, circular points indicate the experimental data values. Every deconvolution was based on the same set of initial parameters with fixed position and fixed FWHM of all Lorentzan peaks and freedom of 5cm^-1^ and 4cm^-1^ respectively. Participation of subpeaks in Amide I deconvoluted spectra over time for **G)** 10% PEG and **J)** 30% PEG systems. Dashed lines are linear fit and circular points represent a peak area for each time of maturation.

This substantial spectral difference between the two regions implies that the crowding agent, PEG, is primarily situated in the continuous phase, with minimal to no presence in the droplet phase. Conversely, the continuous phase demonstrates a relative protein deficiency - the signal emanating from the continuous phase closely mirrors that from the prevalent PEG, with no discernible trace of protein. These observations lead us to deduce that the system exhibits a competitive behavior, characterized by a struggle between the two species to occupy distinct spaces, resulting in fully segregative phase separation. Recently, it has been shown that titration of such “depletant crowders” can be used to quantify polymer-solvent interaction [54].

Since we noticed spatial variations in condensate formation, we surmised that kinetic phenomena play a key role in the phase separation of Nup98. Consequently, we conducted an analysis of the transition of the chemical composition of Nup98 condensates at different PEG concentrations over time. This examination focuses on the chemical evolution of droplets during their maturation process. We instituted a periodic sampling procedure, wherein BCARS measurements were taken at 5 minutes and up to a significant span of 24 hours post the initiation of PS. The addition of PEG instigates pronounced changes within the Raman spectrum, specifically in the CH stretch region of the droplets (**Fig. S2 and S3**), an effect that displays a monotonic positive relationship with the concentration of PEG (**Fig. 1D**). In the incipient phase of the experiment, especially when a 30% PEG concentration is employed, a notable increase in peak intensity at 2990 cm^-1^ and 3065 cm^-1^ was recorded. These are attributed to the glycine residue and the residual asymmetric stretching of the CH_3_ groups, and the breathing mode of the aromatic ring of phenylalanine, respectively. This points towards an increase in hydrophobic intermolecular interactions amongst the FG repeating units of the protein chains and suggests densification of the protein packing within the droplets.

We also noticed a gradual change in the profile of the Amide I peak as the amount of PEG increased: specifically, a decrease in the shoulder at 1645 cm^-1^ over time. To discern the intricacies of the modifications within the secondary structure of Nup98, we performed a spectral deconvolution analysis of the Amide I vibration of Nup98 condensates at different PEG concentrations over time (**Fig. 1E-1J and S4**). In the absence of a crowding agent, a discernible change in the peak shape and a narrowing over time is observed. Initially, a distinct contribution from the alpha-helix and random coil sub-peaks decreases over the course of the experiment and is accompanied by an increase in the contribution from beta-sheet and beta-turn structures (**Fig. 1G**). In the presence of PEG, this effect is noticeably tempered. Samples containing PEG have a similar, Amide I peak profile, even at the early time of maturation, to that of the sample that was not treated with PEG after 24 h of the maturation, albeit with a slightly higher and still rising beta-turn content (**Fig. 1J**). These observations indicate that changes in the secondary structure of Nup98 in condensates occur via beta-sheet formation and that the kinetics of this structural maturation process can be expedited through the introduction of PEG.

The secondary structure of proteins plays a critical role in defining the mechanical properties of protein droplets. The specific folding patterns, such as beta-sheets, dictate the physical strength and elasticity of the droplets, thus influencing their stability and resistance to external stresses [55]-[57]. Additionally, transformations in these structures can lead to alterations in intermolecular interactions that subsequently impact the viscoelasticity of the droplets. Therefore, an understanding of these structural dynamics is fundamental to comprehending the physical behavior of protein droplets under various conditions. To study how the presence of PEG influences the mechanical properties of the droplets during maturation, we performed FRAP over 5 minutes to 24 hours, in the presence of PEG at varying concentrations. We observed a many of amalgamated droplets forming larger spherical condensates during the early stages of the experiment for all samples at 5 minutes post-formation. However, this tendency was notably subdued with increasing PEG concentrations, resulting in droplets of considerably smaller diameters and more irregular shapes (**Fig. 2A** and **Fig. S5**). For our FRAP studies, we matched the same sampling intervals as for BCARS.

**Figure 2.**
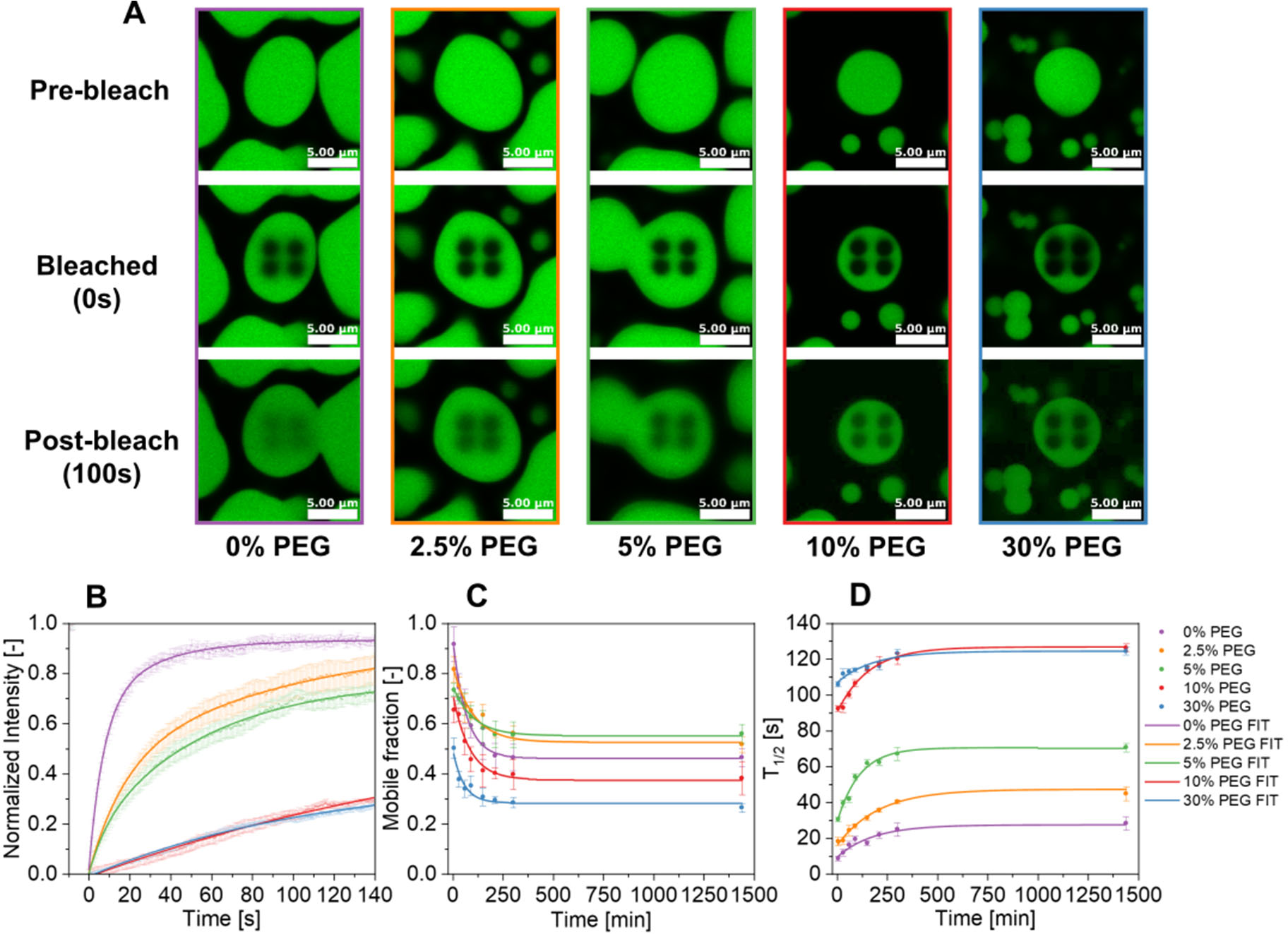
FRAP of Nup98:PEG condensates shows reduced recovery with maturation time. **A)** Representative images from FRAP experiments of 1% Alexa Fluor™ 488-labeled Nup 0.165mM condensates for 0%-30% PEG systems captured 5-30min after initializing LLPS. Each sample was bleached simultaneously in four circular spots with the same diameter of 1.5µm. **B)** Normalized fluorescence intensity of 0%-30% PEG systems over time after bleaching. Presented curves apply to 5min maturated samples. **C)** Dependence of the mobile fraction on maturation time. **D)** Dependence of the half-time fluorescence recovery on droplet time. In graphs B-D thick lines are fit curves, circular points indicate the average values of 4 bleached regions, and error bars show the standard deviation.

Intriguingly, even low mean PEG concentrations (2.5% and 5%) resulted in a noticeable slowing of the fluorescence recovery kinetics, even at 5 min after condensate formation. This deceleration was even stronger by the addition of 10% and 30% PEG (**Fig. 2B**). In the overall analysis, the maturation process prominently exhibited a substantial decline in the mobile fraction (**Fig. 2C**), indicating a reduction in the freely diffusing particles within the droplets. This phenomenon was concurrently paired with a noticeable increase of the half-time recovery (**Fig. 2D**), indicating an extended period required for particle diffusion, as observed from the FRAP experiments. These findings were consistent across all explored concentrations of the crowding agent, suggesting a marked influence on the droplet’s internal dynamics and stability. Dynamics of mobile fraction and half-time recovery changes did not exhibit linear time dependence. Rather, they manifested logarithmic decay and growth, respectively. Increasing the PEG concentration correspondingly amplified the slowing mobility trends, resulting in diminished mobile fractions and protracted half-time recoveries in almost all measured cases. The only exception was a slight increase in the mobile fraction for prolonged maturation times in the presence of lower PEG amounts (2.5% and 5%). Taken together, the structural and physical maturation of Nup98 in the presence of PEG clearly shows that maturation is accelerated at the molecular and microscopic levels. PEG seems to exclude volume artificially concentrating Nup98 and boost protein-protein interaction which results in faster beta-sheet formation and thus reduces liquid-like properties of condensates. The observations herein align with the hypothesis positing that even minimal intermolecular homotypic interactions may be sufficient to establish a percolation pathway. This pathway, in turn, could induce the emergence of dynamically arrested states, culminating in the genesis of gel structures which transform primarily liquid droplets (**Fig. S5**) into gel like aggregates [58], [59].

### PEG crowding drives segregative BSA condensation and accelerates physical maturation

In this section, we study the phase separation and possible maturation of a solution containing PEG and Bovine serum albumin (BSA). BSA is commonly referred to as a protein with a well-defined, stable, secondary structure. This makes it an ideal candidate for a comparison with the behavior of the disordered Nup98 in terms of how PEG crowding affects condensation and maturation in the case of an ordered protein. BSA primarily consists of alpha-helices, which account for up to 67% of its total secondary structure composition [60], [61], resulting in a consistent and reliable Raman signature that serves as a helpful benchmark for spectroscopic analysis. The robustness and temporal stability of BSA, despite fluctuations in environmental factors, underscores its utility in these investigations.

To study phase separation and to discern the impact of PEG on the kinetics of BSA condensate maturation, we used the same protocol as for Nup98. Identical measurement intervals and research techniques were employed, though the PEG concentrations differed. Notably, we observed that BSA does not undergo phase separation if the concentration of PEG is less than ∼10% (w/v) for solutions with an average BSA concentration of 10 μM (**Fig. S6**). This behavior aligns with findings reported by Poudyal *et al.* on 8000 g/mol PEG [34]. We note that BSA alone (in excess of 1 mM), nor PEG alone (even at 40% (w/v)) formed any droplets. Condensation required BSA together with PEG. Furthermore, the noteworthy heterogeneity of the samples warrants mentioning. Unlike Nup98 condensates, BSA droplets were primarily located in the region where the protein solution had been pre-deposited on the coverslip, which hints at a significant diffusion limitation in the mixing of protein with the PEG-containing buffer (**Fig. S6**).

As for Nup98, we analyzed regions of the sample by selecting the locations where both the continuous and droplet phases were clearly observed (**Fig. 3A**). BCARS imaging was initiated in these regions, and we randomly selected a representative number of pixels epitomizing the continuous and droplets phases for spectral analysis (**Fig. 3B**). The acquired spectra from these areas were compared with the spectra of solutions of the pure BSA and PEG (**Fig. 3C**). The continuous phase exhibited a Raman-like spectrum almost identical to that of the PEG solution, with common characteristic peaks in both the fingerprint and CH stretch regions. The droplet phase provides more intriguing insights. The fingerprint region contains peaks from both reference spectra, particularly apparent in the peak at 1280 cm^-1^ and the double peak at ∼1455 cm^-1^. The upper part can be attributed to PEG, whereas the lower one (closer to 1440 cm^-1^) is associated with BSA, related to various deformations of the CH_2_ groups [49]. This shows that, in contrast to Nup98:PEG, the BSA-rich droplets contain a significant fraction of PEG and that the spectrum of the droplet phase in the CH stretch region assumes a unique shape that does not replicate either of the reference spectra.

**Figure 3.**
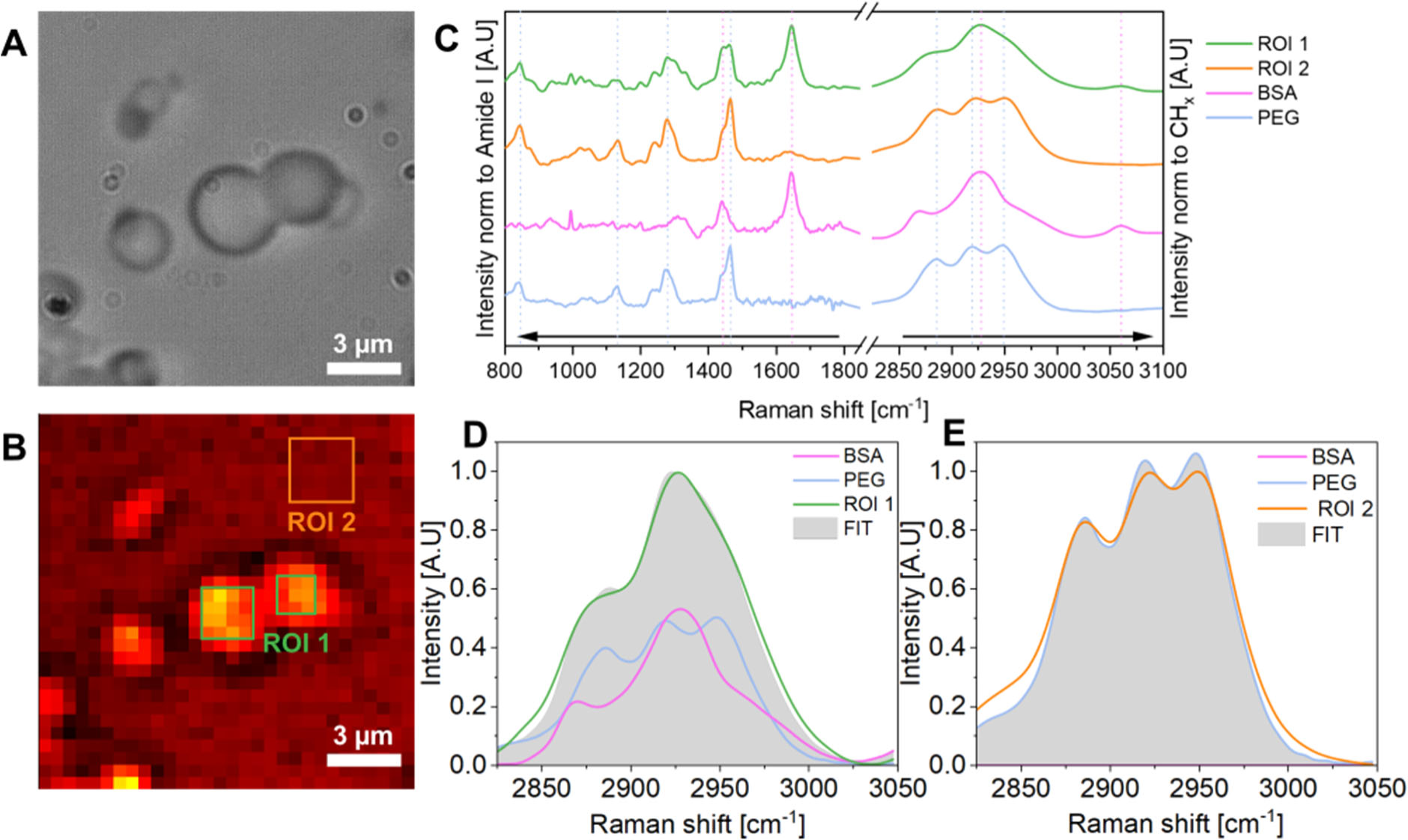
Molecular microscopy of BSA:PEG condensate aging. In situ **A)** BF image and **B)** BCARS map imaging of the same BSA condensates formed by mixing 1:10 BSA 0.125mM with PEG 4kDa 75mM (30% m/v). Both images were captured 24h after initiation of LLPS. BCARS map shows averaged CH stretch intensity and is integrated over 2820 - 3020cm^-1^ with a pixel size of 0.5×0.5µm. On the BCARS map, ROI 1 marked green, represents a selected part of the droplet phase, and ROI 2 marked orange, represents a selected part of the continuous phase of a sample. **C)** Normalized fingerprint and CH stretch spectra comparison of ROI 1 and ROI 2 to 4mM BSA and 75mM PEG reference samples. **D)** ROI 1 **E)** ROI 2 CH stretch spectral fitting based on BSA and PEG reference samples. Fit is a grey area, and it is a linear sum of BSA and PEG contributions.

In a more detailed quantitative analysis, we conducted a fit based on the aforementioned reference spectra. As suggested above, the droplet phase appears to comprise both PEG and BSA (**Fig. 3D**). Conversely, the continuous phase is almost devoid of protein (**Fig. 3E**). From the fits, we estimated the concentrations of BSA and PEG in both phases (**Table 1**). It is evident that the concentration of PEG in the droplet phase is significant, with its value non-linearly increasing with the mean PEG concentration, but never exceeding it. On the other hand, BSA is concentrated in the droplets by 70-190 times the mean value, depending on the mean PEG concentration. The BSA concentration in the continuous phase is below the detection limit and displays a dominance of the crowding agent. In other words, the phase separation of BSA:PEG:buffer is segregative as well, though not as pronounced as for Nup98:PEG:buffer. Since the mean BSA concentration is very low (see **Table 1**), the total volume of the dispersed phase is much smaller than that of the continuous phase, implying that the PEG concentration in the dilute phase should be close to its mean value. **Table 1** demonstrates that our spectral quantification method aptly substantiates this, presenting a slight underestimation, which modestly amplifies with the PEG concentration. Intriguingly, our approach to determining the anticipated PEG concentration in the continuous phase employs a strategy founded on the principle of mass conservation, ensuring that the total amount of PEG remains constant within the system even as it partitions between phases. The process through which we calculate the expected PEG concentration in the continuous phase is further elucidated in the **SI, Section S7**.

**Table 1.** Summary of BSA and PEG concentrations for different systems. *Indicates directly calculated from BCARS spectra

| PEG content | Component | Before mixing<br>[mg·ml<sup>-1</sup>] | Theoretical dilution<br>[mg·ml<sup>-1</sup>] | Droplet phase*<br>[mg·ml<sup>-1</sup>] | Continuous phase*<br>[mg·ml<sup>-1</sup>] | Expected continuous phase<br>[mg·ml<sup>-1</sup>] |
| --- | --- | --- | --- | --- | --- | --- |
| 10% PEG | PEG | 100 | 90.90 | 51.72 | 91.26 | 91.47 |
|  | BSA | 8.25 | 0.77 | 53.59 | N.A. | 0 |
| 20% PEG | PEG | 200 | 181.82 | 88.17 | 167.92 | 182.66 |
|  | BSA | 8.25 | 0.77 | 83.95 | N.A. | 0 |
| 30% PEG | PEG | 300 | 272.73 | 108.90 | 247.82 | 273.56 |
|  | BSA | 8.25 | 0.77 | 148.30 | N.A. | 0 |

The fact that, in contrast to the Nup98 solutions, the PEG concentration in the dispersed BSA-rich phase gradually increases with the mean value, provides the opportunity to elucidate the interaction network that determines the phase diagram. We accomplish this by fitting the experimental binodal concentrations against a free energy model that combines Flory-Huggins theory [67] with statistical associating fluid theory (SAFT) [68]. This type of hybrid model has shown to be an excellent choice for interpreting and predicting biomolecular phase behavior [11], [69], [54] as it discriminates between non-specific interactions, captured in a general way by the term “solvation”, and specific association between defined sites located along the polymer backbone. In defining the model, we consider the solution of BSA, PEG and buffer as a ternary mixture, of which the species are respectively referred to as components A, B, and S (the latter being from “solvent”). The model considers the propensity of BSA to non-covalently dimerize [70] and to some extent form higher-order complexes [71], though without considering the exact mode of association. We define the following binding equilibrium (**Fig. 4A**).

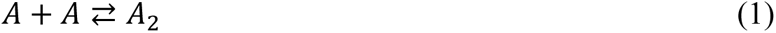

**Figure 4.**
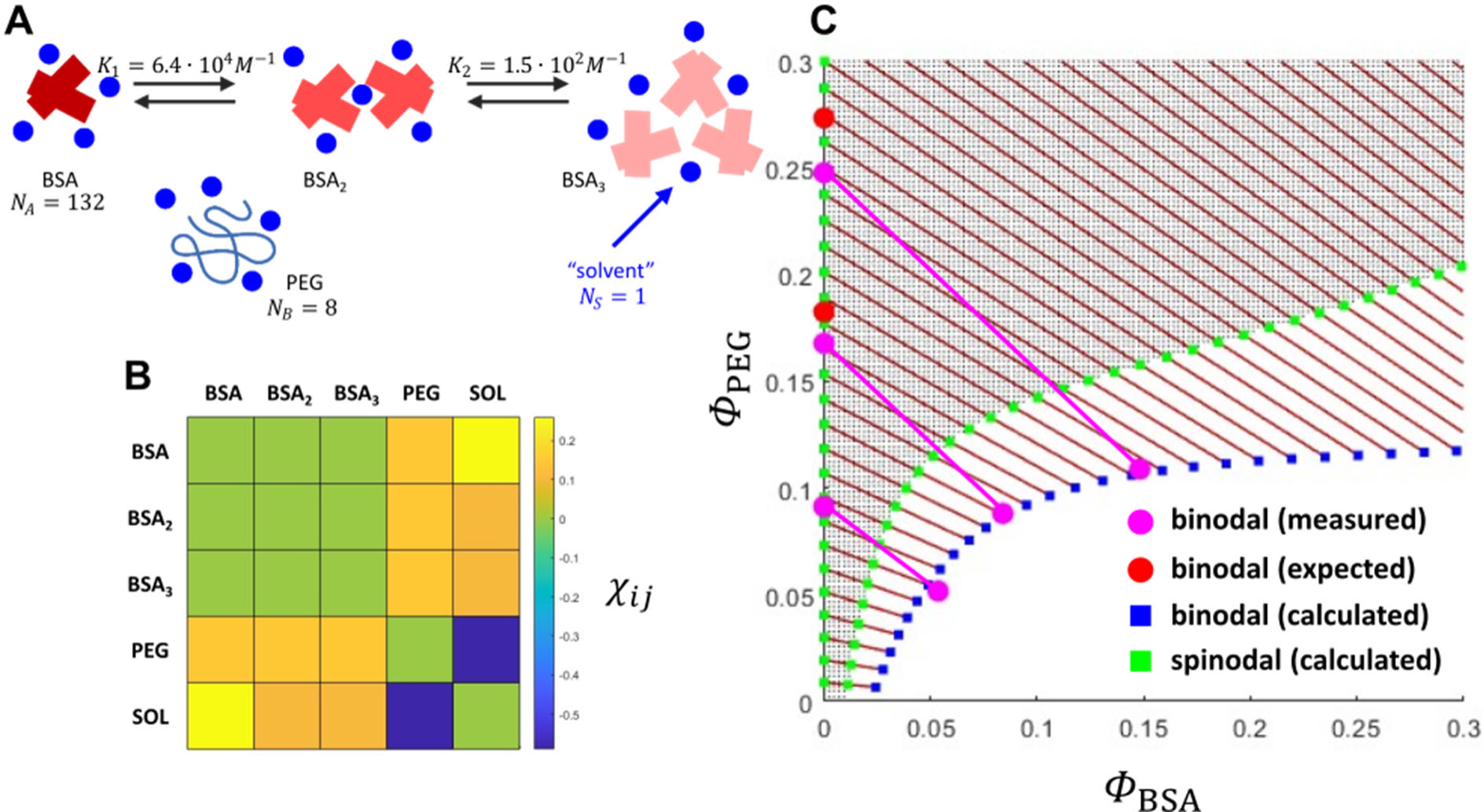
Phase field modeling predicts BSA:PEG segregative phase separation **A)** Schematic overview of the individual species presumed present in the binary solution BSA:PEG:solvent, together with association constants and relative molecular sizes obtained from the. fitting **B)** matrix of binary interaction parameters, evaluated from fitting **C)** experimental and fitted (calculated) phase diagram. The red dots on the protein-devoid binodal branch represent expected data, calculated on the basis of mass conservation, assuming the concentrations in the protein-rich phase to be correct. The deviation from the measured point seems to increase with PEG concentration.

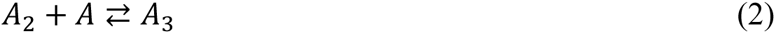

, characterized by the association constants:

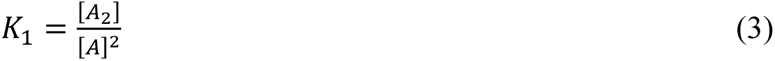

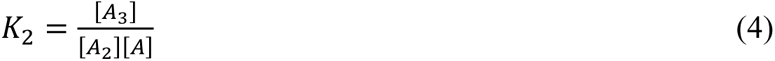

, where the square brackets denote molar concentrations.

The dimensionless free energy density of the mixture comprises three contributions: one from translational entropy, one from non-specific interaction and one from non-covalent binding:

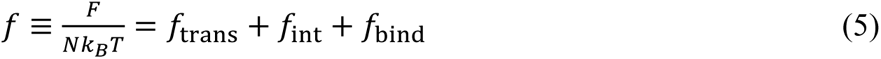

Here, *F* is the total free energy, *k_B_T* the thermal energy and *N* the total number of sites of an imaginary molecular lattice, onto which we map the mixture [67]. The first and second term on the RHS of Equation (5) are given according to Flory-Huggins theory:

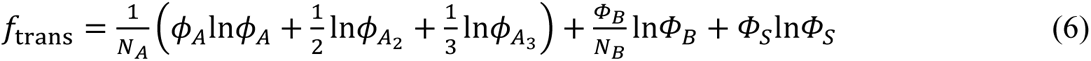

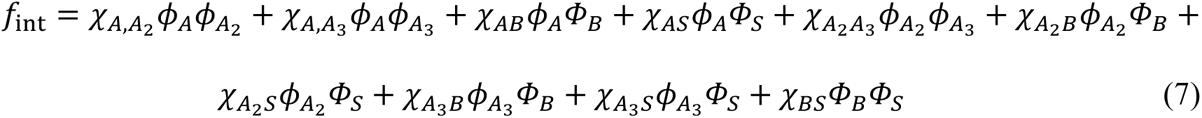

Here, *Φ*_*A*_, *Φ*_*A*2_ and *Φ*_*A*3_ are the equilibrium volume fractions of BSA monomers, dimers and trimers, with the total BSA volume fraction given by: *Φ*_*A*_ = *Φ*_*A*_ + *Φ*_*A*2_ + *Φ*_*A*3_. The parameters *N*_*i*_ denote the effective molecular sizes in terms of lattice sites, *Φ*_*B*_ and *Φ*_*S*_ = 1 - *Φ*_*A*_ - *Φ*_*B*_ are the volume fractions of PEG and solvent and *χ*_*ij*_ are Flory parameters. The contribution due to the self-association of BSA is given by

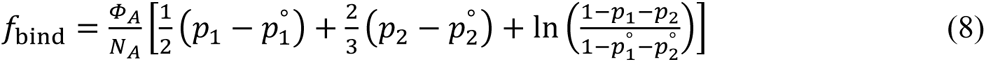

with *p*_1_ and *p*_2_ the fractions of BSA accommodated in dimers and trimers. The superscript ° refers to the pure state. The magnitude of *p*_1_ and *p*_2_ follows from the equilibrium condition: 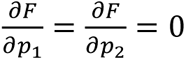 and the condition that the molecular volume ratio of monomer to dimer to trimer is ∼1/2/3. These constraints give rise to the following conditions:

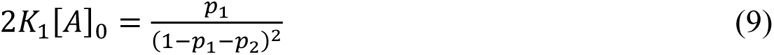

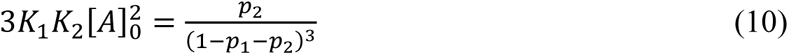

Here, [*A*]_0_ is the total molar concentration of BSA. We refer to the **SI, Section S7** for the derivation of **Eq. 8, 9, 10.**

The fit of the model against the experimental data is represented by the calculated phase diagram in **Fig. 4C**, obtained for effective sizes *N*_*i*_ and association constants *K*_1_ and *K*_2_ as specified in **Fig. 4A**, as well as the Flory parameter interaction matrix (*χ*_*ij*_) given in **Fig. 4B**. The binodal, *i.e.,* the actual fit, was calculated by equalizing the chemical potentials, as well as the osmotic pressure in the coexisting phases, whereas the spinodal was obtained by the condition det(**H**) = 0, with **H** the Hessian matrix of second derivatives of *f* with respect to composition. The fitting was subject to seven constraints to limit the number of free-floating variables and eliminate non-uniqueness. In **Table 2** we list these, together with a short motivation. Note that constraints 4, 5, and 6 reduce the number of relevant Flory parameters from ten to four.

**Table 2.**
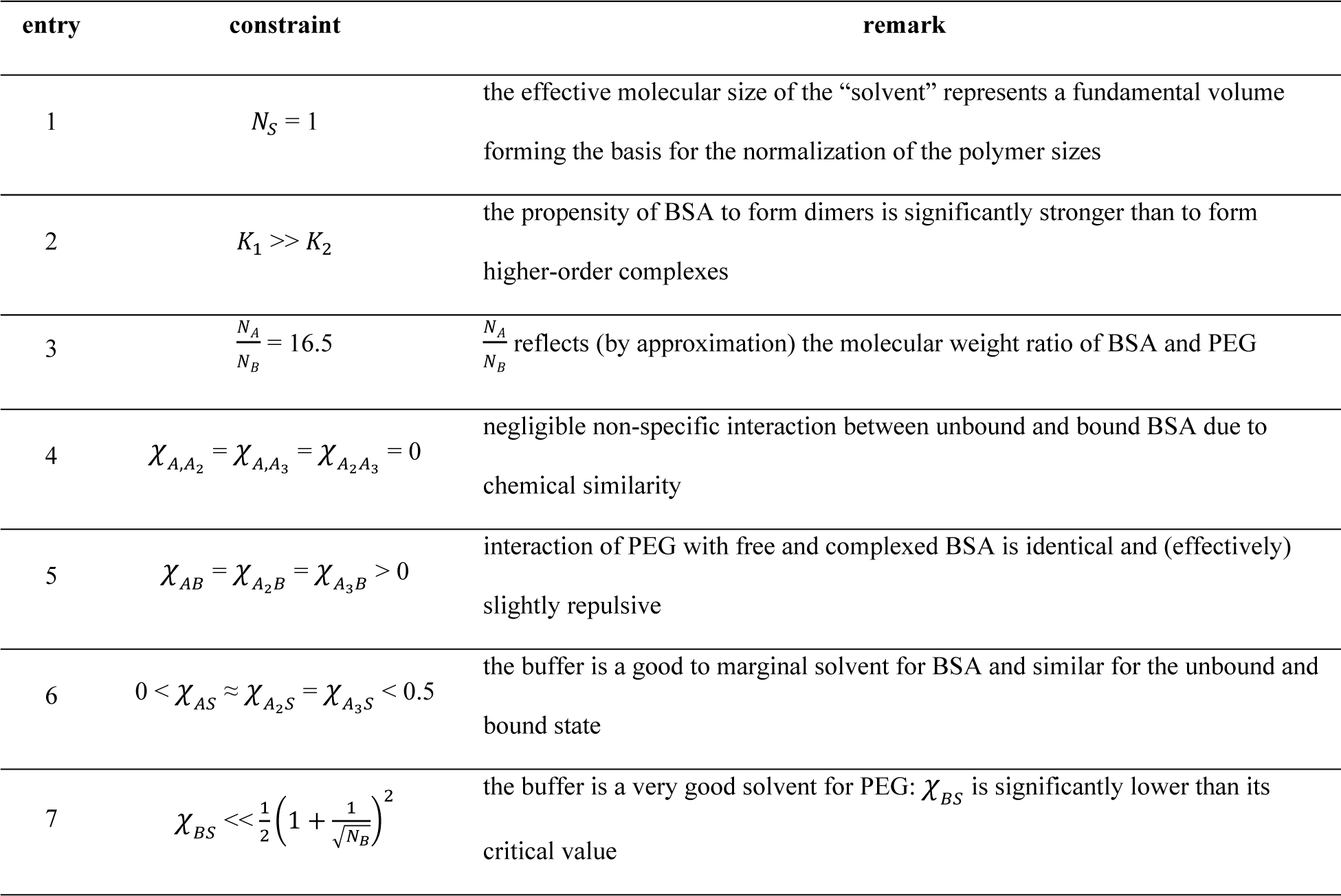
Constraints for fitting the BSA:PEG:buffer phase diagram.

**Figure 4** shows that the calculated binodal excellently fits the experimental results. Of the latter, the pink and red symbols mark measured and expected data (see **Table 1** and above). There is a small discrepancy in the tie-line angle, but the model reproduces the segregative phase separation and confirms the presence of a significant fraction of PEG in the BSA-rich phase. Our value for the dimerization constant *K*_1_ agrees well with the value of ∼10^5^ M^-1^, reported by Levi *et al.* [70] and significantly exceeds *K*_2_. This shows that under our experimental conditions, dimerization of BSA dominates the formation of higher-order complexes. We consider the listed *K*_2_ a maximum, since values down by one order of magnitude did not affect the quality of the fit. This result is hence in agreement with the observation that near room temperature aggregation of BSA is negligible [71]. Regarding the non-specific interactions, it is interesting to note that *χ*_*AS*_ somewhat exceeds *χ*_*A*2*S*_ and *χ*_*A*3*S*_, implying the BSA complexes to be slightly better solvated than unbound BSA. Finally, the fact that *χ*_*BS*_ turns out negative is consistent with the notion that low molecular weight PEG is very hygroscopic and even deliquescent [72], [73], [74] (though its hydrophilicity decreases with increasing molecular weight). This result suggests that the “crowding action” of this low molecular weight PEG mainly stems from its osmolyte nature.

A notable observation is that in the solutions containing 10% and 30% PEG the Amide I peak remains largely unchanged throughout the 24 h experiment, with major contributions from alpha helices and random coils. This observation is consistent with the well-structured BSA, which is characterized by an alpha-helical dominance (**Fig. S8, S9, and S10**). However, in the 10% PEG sample, we discern a progressive alteration in the ratio of random coils to alpha-helix over time (**Fig. S8 and S10**). Similarly, a subtle change in the spectral shape can also be seen for the CH stretch region for 10% PEG in the emergence of a shoulder at 2950 cm^-1^, which might suggest a slow change in PEG:BSA ratio inside condensates (**Fig. S8, S9**). In the case of 30% PEG, neither significant changes in Amide I nor CH stretch regions are observed during the 24 h experiment (**Fig. S8, S9**). This behavior elucidates that, in the frame of maturation within the BSA-PEG system, any slight alterations in the secondary structure, particularly as witnessed in the 10% PEG system, are not a predominant effect of maturation. Specifically, variations within the secondary structures, such as those within the alpha-helix, may be attributed to mere fluctuations, especially in instances where the protein is not sufficiently crowded, as within the 10% PEG condition. Furthermore, higher concentrations of PEG confer stability to the system, enhancing this cooperative dynamic and ensuring that the process is not significantly influenced by minor perturbations in protein structure but is governed by the synergistic participation of both PEG and BSA in the droplet phase.

The composition of biomolecular condensates can profoundly impact their mechanical properties, which is a crucial factor in their purported role in biological processes. *In vitro*, the presence of a crowding agent inside the droplets can modulate their dynamics [62], [63]. When a crowding agent permeates the droplet, it can influence the viscosity and elasticity of the droplet, hence affecting its physical behavior [64]–[66]. In addition, the molecules of the crowding agent can contribute to a decrease in the diffusivity of proteins within the droplet. This transition can also influence the interactions between the droplets and their environment. Consequently, comprehending how the presence of crowding agents modulates attributes such as liquidity and viscosity within the droplets provides vital insights into the inherent characteristics and functionalities of these microscopic entities.

To explore changes in droplets’ mechanical properties and protein chains’ diffusability inside condensates, with a particular emphasis on the dynamic properties studied over time as a function of PEG content, we used FRAP (Fig. 5A). In accordance with the established protocol, the time intervals for these measurements were maintained identical as for Nup98 based systems. However, in contrast to the Nup98 systems, it was not possible to fully deplete the intensity of the bleached regions in the droplets to near-zero levels. This observation suggests that the recovery of the system might occur on a timescale shorter than our frame capture rate, indicating that the recovery kinetics we are measuring could be an overestimation, particularly in terms of the mobile fraction Hence, for this system we refer to this measure as the “apparent mobile fraction”.

**Figure 5.**
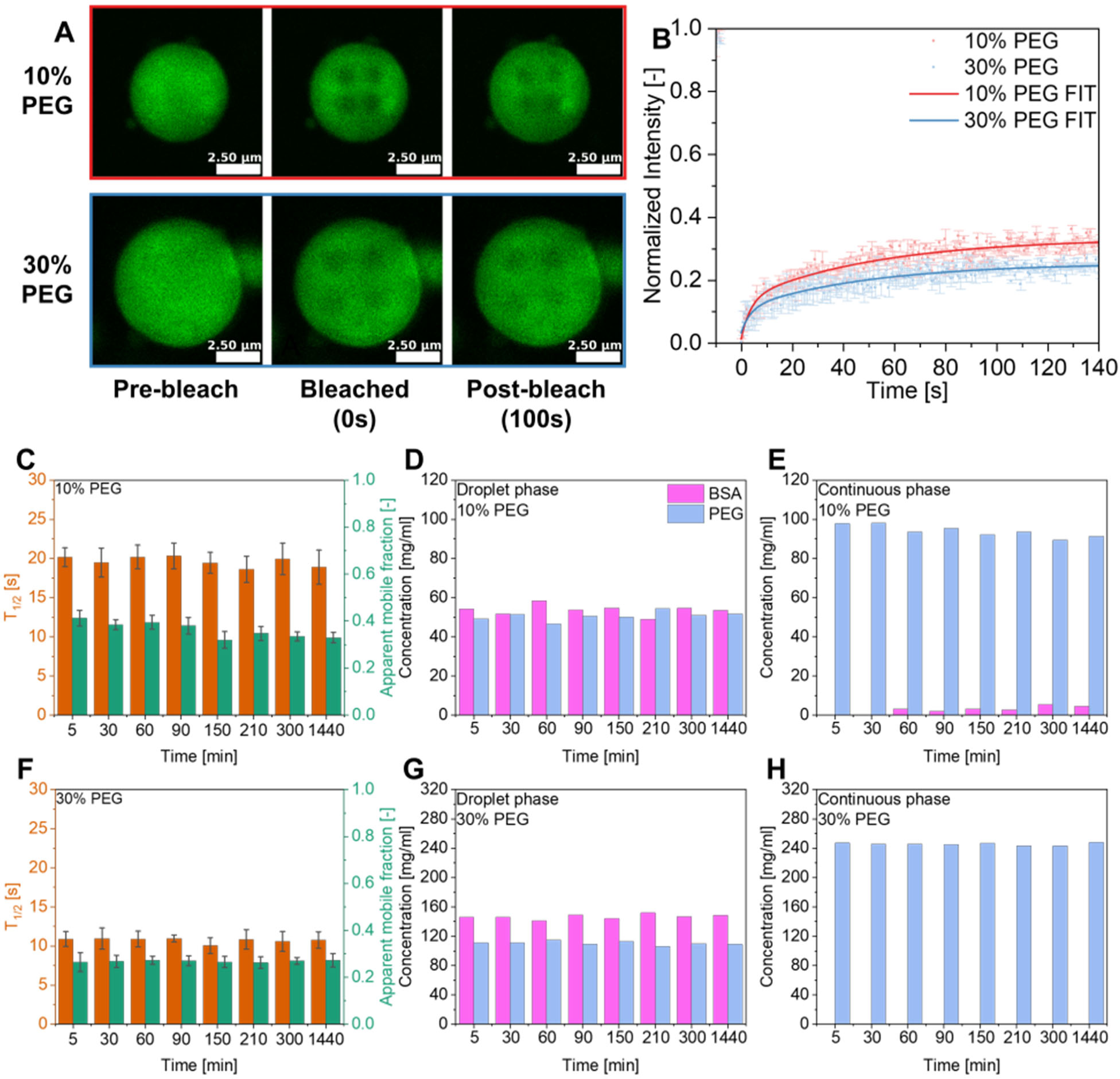
FRAP and concentration analysis of BSA:PEG condensate aging. **A)** Representative images from FRAP experiments of 1% FITC-labeled BSA 0.125mM condensates for 10% PEG and 30% PEG systems captured 5-30min after initializing LLPS. Each sample was bleached simultaneously in four circular spots with the same diameter of 0.8µm (10% PEG) and 1.0 µm (30% PEG). **B)** Normalized fluorescence intensity of 10% PEG and 30% PEG systems over time after bleaching. Presented curves apply to 24h maturated samples. Thick lines are fit curves, circular points indicate the average values of 4 bleached regions, and error bars show the standard deviation. Bar graphs of half-time fluorescence recovery and apparent mobility fraction for **C)** 10% PEG and **F)** 30% PEG systems for different maturation times. Error bars are standard deviations. Concentrations of BSA and PEG over time in the droplet phase for **D)** 10% PEG and **G)** 30% PEG, and continuous phase for **E)** 10% PEG, and **H)** 30% PEG systems.

From the beginning of the experiment both 10% and 30% PEG concentrations exhibit analogous kinetics in the fluorescence recovery process, as evidenced by the similar shapes of their fluorescence recovery curves (Fig. 5B). Moreover, we noticed a consistent stability of both the apparent mobile fraction and the half-time of recovery throughout the duration of the 24-hour experiment (Fig. 5C and 5F) for both PEG concentrations. However, when the PEG concentration was at 10%, we identified marginally more pronounced fluctuations in the half-time of recovery and apparent mobility fraction values. These observations are qualitatively consistent with the Raman data collected on droplet composition for the 10% PEG system, where subtle changes in the Amide I and CH stretch spectral regions were also observed in the experiment (Fig. 5D). Interestingly, a noticeable rise in BSA concentration was observed in the continuous phase for 10% PEG, where the initial concentration was below the detection limit at the beginning of the experiment. After approximately 60 minutes, the concentration rose significantly and exceeded the theoretical mixing concentration (Fig. 5E). This prompts us to consider that a 10% PEG concentration may not be adequately sufficient to thoroughly crowd BSA. Modeling intimates that a reduction in crowding agent concentration could steer results toward approaching a possible critical point where both metastable and stable separation regions are separated by slight fraction values. Consequently, minor variations in local concentrations might be a driving force for all observed fluctuations in both Raman and FRAP studies. However, when considering cases where the PEG concentration was 30%, we did not observe a similar change in protein or PEG concentration in either phase (Fig. 5G and 5H). Conversely, at a PEG concentration of 30%, the phase separation process is spontaneous, and a well-defined demixed state is preferable.

## DISCUSSION AND SUMMARY

In this work, we aimed to unravel the intricacies of how PEG interacts with protein droplets, discovering that its influence is multifaceted and varies depending on the particular protein in question. A major observation is that the aging process of protein droplets under the influence of PEG is not uniform across different proteins. When examining Nup98, a disordered protein with many FG tandem repeats, we found that PEG considerably accelerates the chemical and structural changes in the protein when condensed. The secondary structure of Nup98 exhibits maturation via the formation of beta-sheets in droplets, which is accelerated when PEG is present. We postulate this is because PEG facilitates a denser packing of protein molecules inside the droplets, thereby enhancing their protein-protein sequence-specific, sticker-like, hydrophobic interactions and effectively speeding up the kinetics of the maturation process. As established by others, these specific interactions are not essential for phase separation [34]. Yet, when a crowding agent is introduced, it not only increases the significance of these interactions but may also alter the kinetics associated with changes in viscoelastic and aging properties [75]. Our results with PEG and Nup98 also offer robust support for the hypothesis that even limited, but specific, intermolecular Nup98-Nup98 interactions boosted by the crowder can stimulate the formation of a percolated network of Nup98 chains. The establishment of such a network is understood to be a pivotal step toward the manifestation of dynamically arrested states, characterized by their significant loss of molecular mobility. This dynamic arrest, once initiated, appears to be a mechanistic pathway leading to the formation of a gel-like structure and delineates a transition from a liquid-like state to a solid-like behavior, which is a hallmark of gelation phenomena observed in various protein and RNA systems [58], [59].

Despite how PEG accelerates Nup98 droplet aging, the PEG exhibits a sufficiently strong net repulsive interaction with the Nup98 as evidenced by fully segregative phase separation; this means that the PEG acts as a true depletant in this case. This behavior aligns closely with the concept of an inert crowding agent, wherein the agent exerts a passive influence on the protein. The kinetics of maturation vary based on the amount of crowding agent added to the system. Our observations indicate that phase separation in the ternary mixture tends to expedite the overall reduction of the mixture’s free energy in comparison to binary systems. Introducing even a small quantity of PEG, leads to more rapid system stabilization and faster maturation compared to no PEG. Conversely, larger PEG concentrations rapidly solidify as evidenced by limited fusion, reduced mobile fraction, and long half-time recovery when compared to similar maturation times for droplets with less or no PEG. This suggests the presence of two distinct time scales: the first for forming sufficient homotypic interactions to induce phase separation and yield liquid droplets, and a second associated with the gradual shift from liquid condensates to gel-like aggregates. Furthermore, the crowding agent-dense environment may hasten the beta sheet formation, which could facilitate protein aggregation and amyloid formation. Consequently, this could influence the biological processes triggered by protein piling [76], [77].

In case of BSA, we observe a distinct interaction with the same PEG crowder. BSA, a structured protein, does not phase-separate on its own, even at concentrations exceeding 100 mg/mL. Yet, adding PEG to the mixture (as low as 5%) alters this behavior such that BSA phase separates at mixture concentrations below 1 mg/mL. Unlike Nup98, BSA droplets contained a significant amount of PEG, showing the net interaction between the polymers to be less repulsive than in case of PEG:Nup98. Prior studies have shown a similar pattern, where a high concentration of crowding agent is present inside the droplet phase for proteins [12], [78]. Again, in contrast to Nup98:PEG, we observed no changes in secondary structure for the BSA:PEG system and only minimal changes in molecular mobility from FRAP measurements over the 24-hour measurement period. Our calculations show that a significant contribution to the segregative phase separation between PEG and BSA is the hydrophilicity of the PEG: it effectively enhances the driving force for phase separation by competing for the solvating environment, thereby favoring homotypic interactions between individual BSA molecules. We therefore conclude that (low molecular weight) PEG acts as an osmolyte, similar to, for instance, a hygroscopic salt or highly polar molecules, such as trimethylamine N-oxide (TMAO).

Consistent with a non-repulsive, segregative phase separation, for the 10% PEG:BSA, system we detect a small amount of BSA in the surrounding phase, which increases with time. This is consistent with the proximity of or approach to a critical point, for which the protein-devoid branch of the binodal moves away from the composition axis. Nevertheless, any changes in droplet chemistry or mechanics are minimal during the entire examination period, and such changes are even smaller for high PEG concentration. Consequently, our primary observation is rapid separation of the protein from the solution, devoid of any significant molecular or mechanical maturation of the protein in the droplets within our 24-hour examination period. We cannot, however, exclude that BSA droplet maturation occurred on a timescale faster than our first measurements (5 min post droplet formation) through the second structure of BSA remains essentially unchanged relative to a pure BSA solution.

Our research shows that the use of a suitable crowding agent can trigger phase separation across diverse protein families. Nevertheless, the exact mechanism of crowding varies depending on the properties of the specific protein exposed to the crowding agent. The mechanism of stimulating phase separation of the protein may occur based on: 1) expulsion of the protein from the solution into a dense phase via a net repulsive interaction and 2) an osmolyte action, wherein the crowder competes with the protein for solvation by the aqueous environment and thus promotes protein-protein interaction without being repulsive. A third mechanism, which we do not observe, is a possible attractive interaction between the protein and crowder to induce an associative phase separation. Thus, crowders in protein phase separation encompass rich behaviors that promote different phenomena from protein aging to droplet formation, depending on the specific proteins involved.

## ACKNOWLEDGEMENTS

S.H.P., E.A.L and J.J.M acknowledge support from the Deutsche Forschungsgemeinschaft (DFG) SPP 2191 Molecular Mechanism of functional Phase separation (Project Nr: 402723784). S.H.P. acknowledges support from the DFG #PA2526/3-1/2 and Welch Foundation F-2008-20220331 and National Science Foundation (#2146549). J.J.M. acknowledges the DFG #MI2212/1-1/2. M.Y. was funded by the MSCA Individual Fellowship (TFNUP 89410) and a Humboldt Research Fellowship for Postdoctoral Researchers. T.S. was funded by the EMBO Postdoctoral Fellowship (ALTF 1020-2020). The content is solely the responsibility of the authors and does not necessarily represent the official views of the funding agencies.

## AUTHOR CONTRIBUTIONS

M.B., J.J.M., and S.H.P. designed and conceived the study. M.B. and P.A.G. performed and analyzed the CARS experiments with support from S.H.P. M.B. performed and analyzed FRAP studies. J.J.M. performed the model calculations and analyzed the resulting data. T.S. M.Y., and E.A.L. analytical reagents and protocols. M.B., J.J.M, and S.H.P. wrote the manuscript with comments from all authors.

## Notes

### Competing Interest Statement

The authors have declared no competing interest.

