## Supplemental figures for "Protein-specific crowding accelerates aging in phase-separated droplets"

SUPPORTING INFORMATION

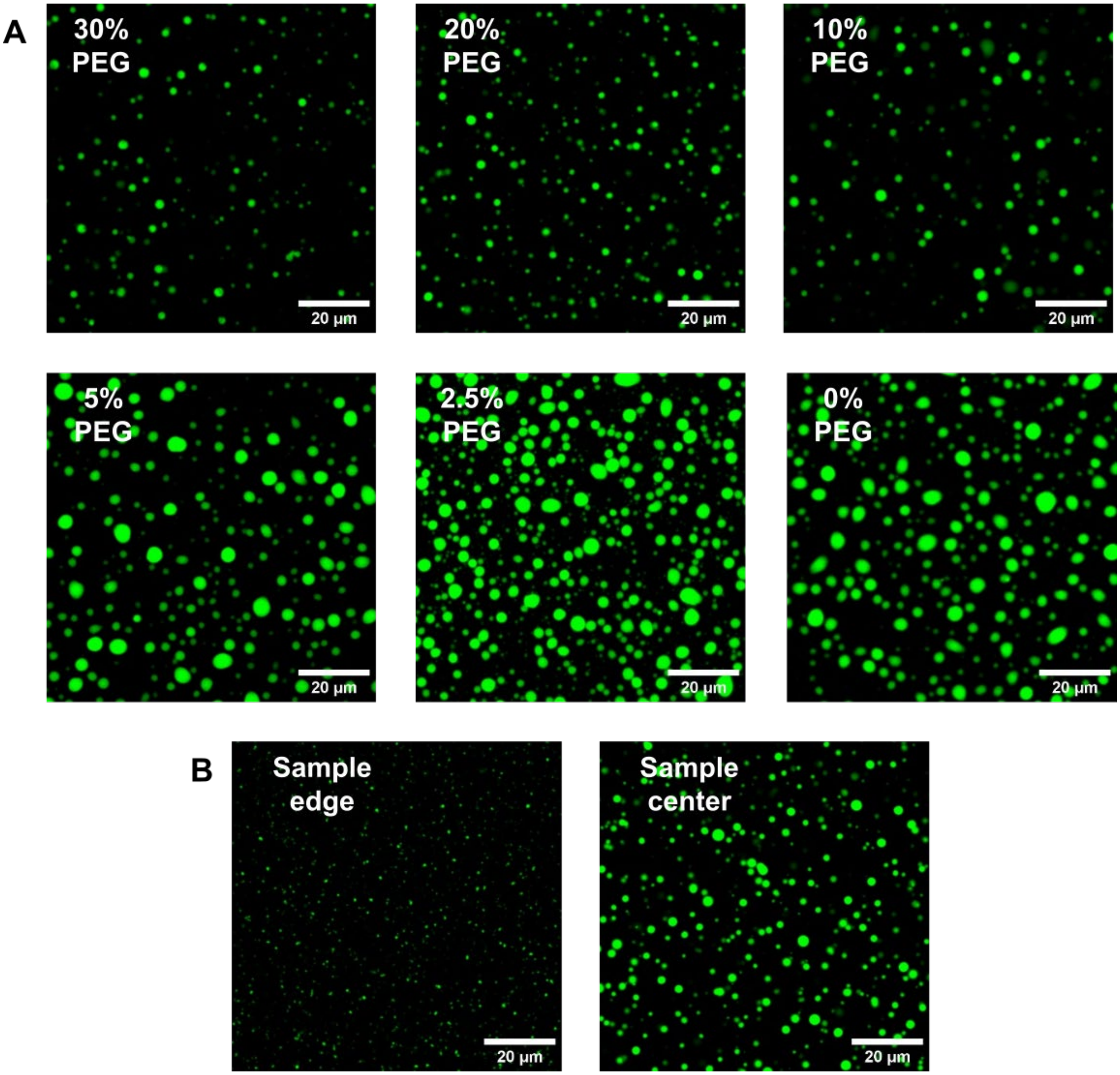


***Figure S1***. *Confocal laser scanning microscopy (CLSM) images of 1% Alexa Fluor™ 488-labeled Nup98 (total concentration 0.165mM ) in Nup98:PEG condensates.* ***A****) 0%-30% Nup98:PEG systems captured 5 min after initializing LLPS in the center of samples,* ***B****) 30% PEG system captured 5 min after initializing LLPS showing different sample positions.*

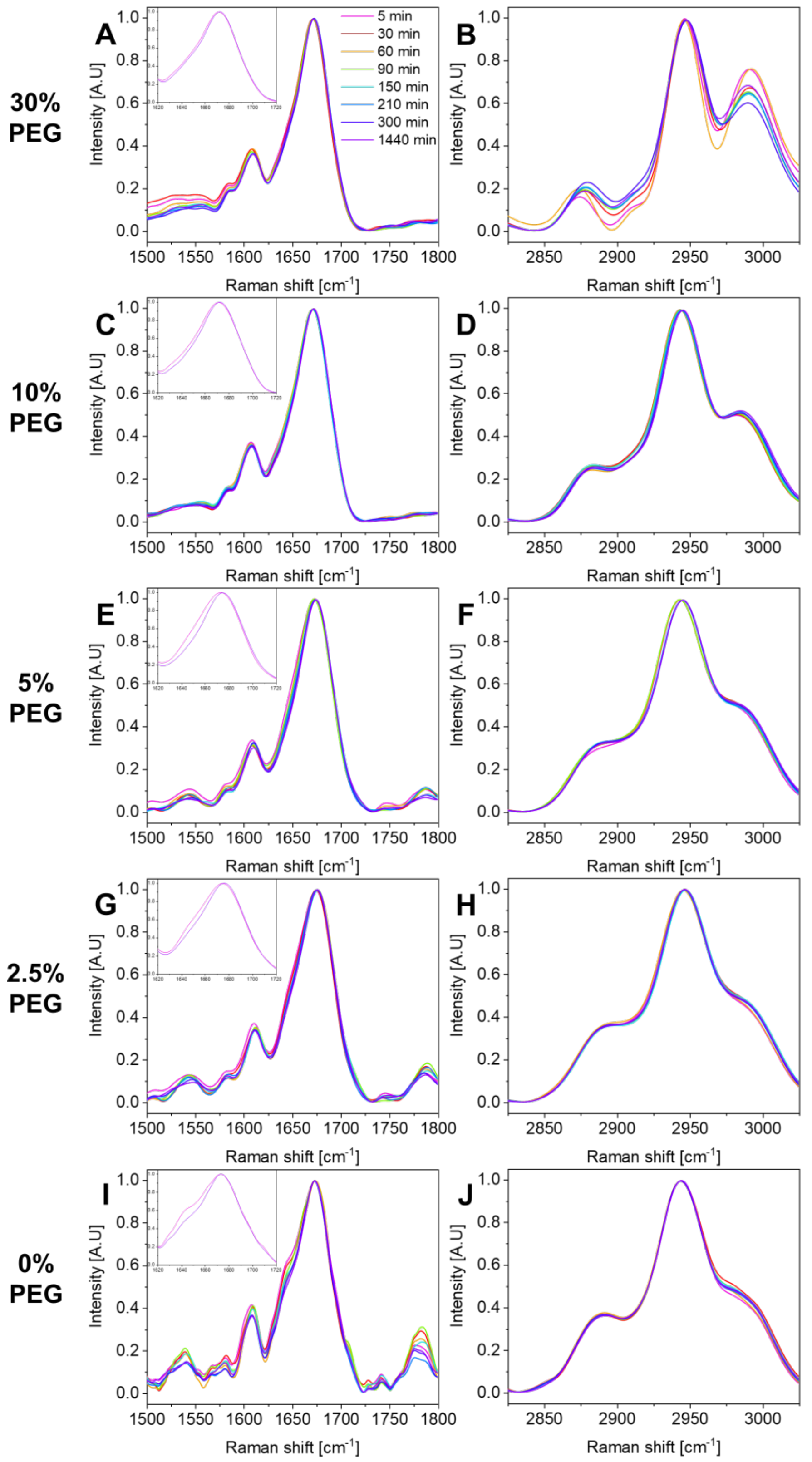


***Figure S2***. *Normalized BCARS fingerprint (**A**, **C**, **E**, **G,** and **I**) and $CH_x$ (**B**, **D**, **F**, **H** and **J**) spectra comparison for droplet phase of Nup98:PEG system at various PEG concentrations over progressive maturation time.*

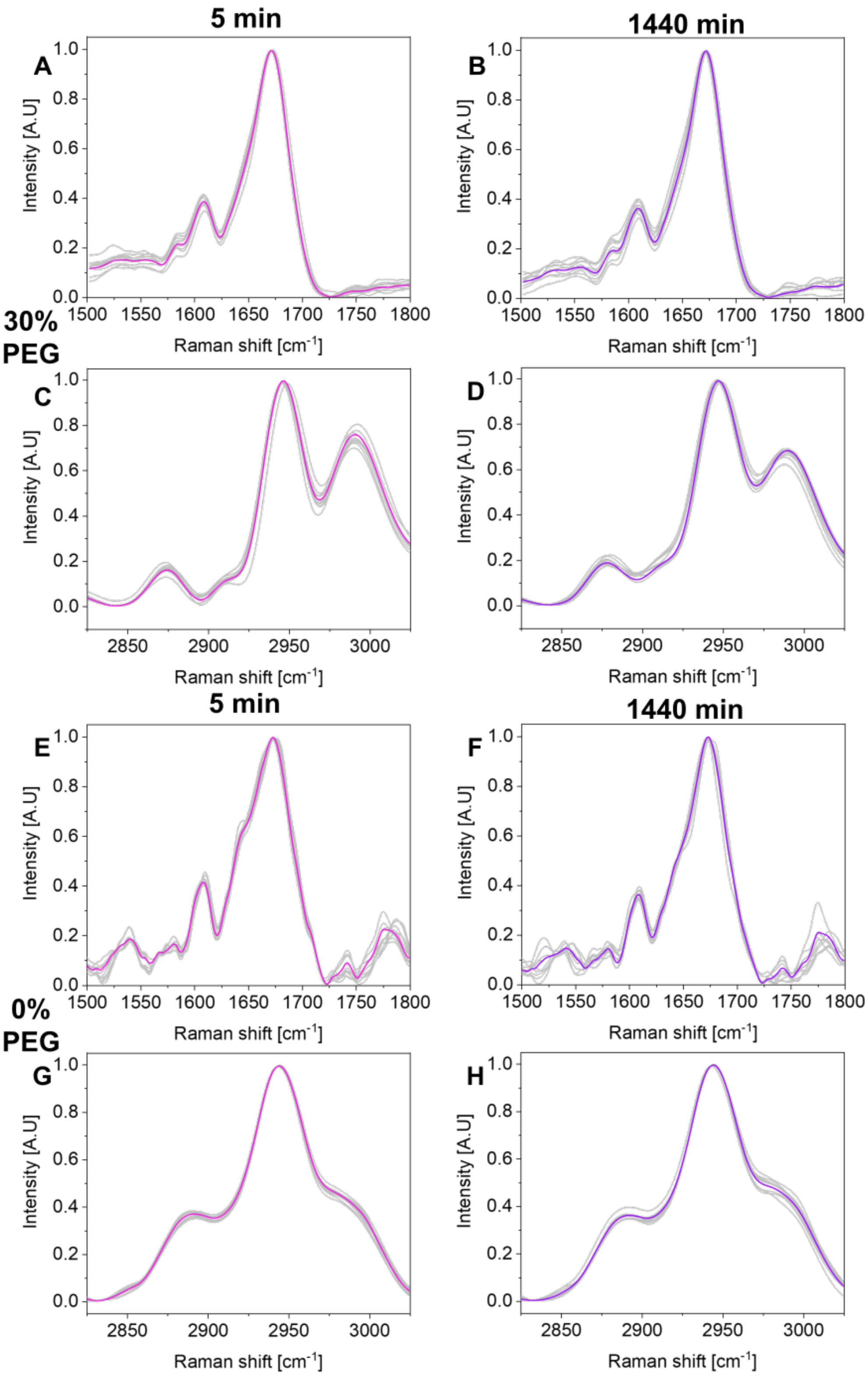


***Figure S3***. *Normalized BCARS fingerprint (**A**, **B**, **E**, **F**) and $CH_x$ (**C**, **D**, **G**, **H**) spectra comparison for droplet phase of Nup98:PEG system for 30% PEG (A, B, C, D) and 10% PEG (E, F, G, H) at 5 min (A, C, E, G) and 1440 min (B, D, F, H). Color lines are averaged data. Grey curves represent 10 exemplary data curves each integrated from >5 individual pixels.*

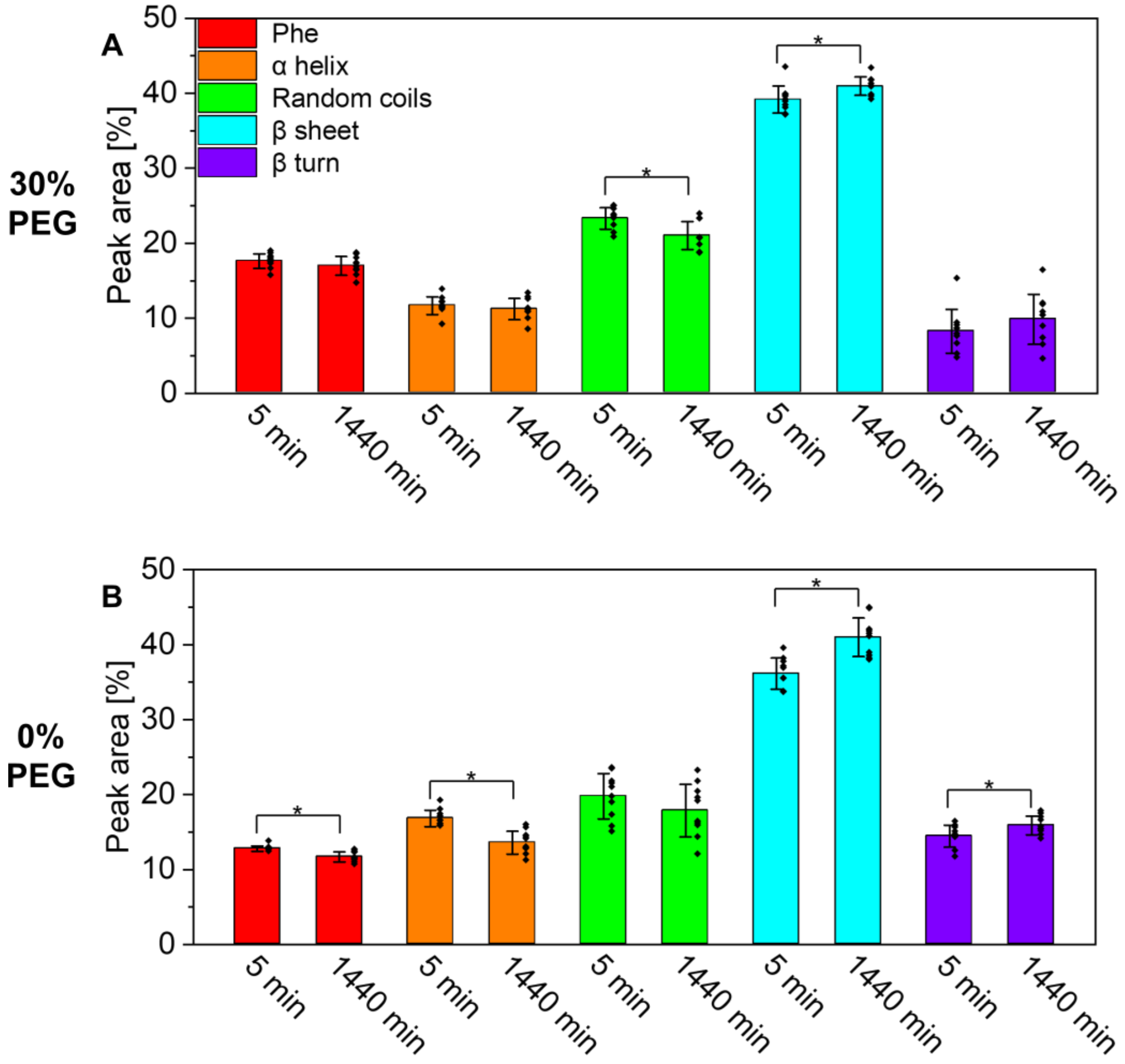


***Figure S4***. *Contribution of subpeaks in Amide I deconvoluted spectra of Nup98:PEG condensate systems* ***A****) 30% PEG and* ***B****) 0% PEG at 5 min and 1440 min. Sample population matches those in Figure S3; n= 10; error bars are std; asterisks show statistical difference of mean values (one-way ANOVA p<0.05), black dots are unique data points.*

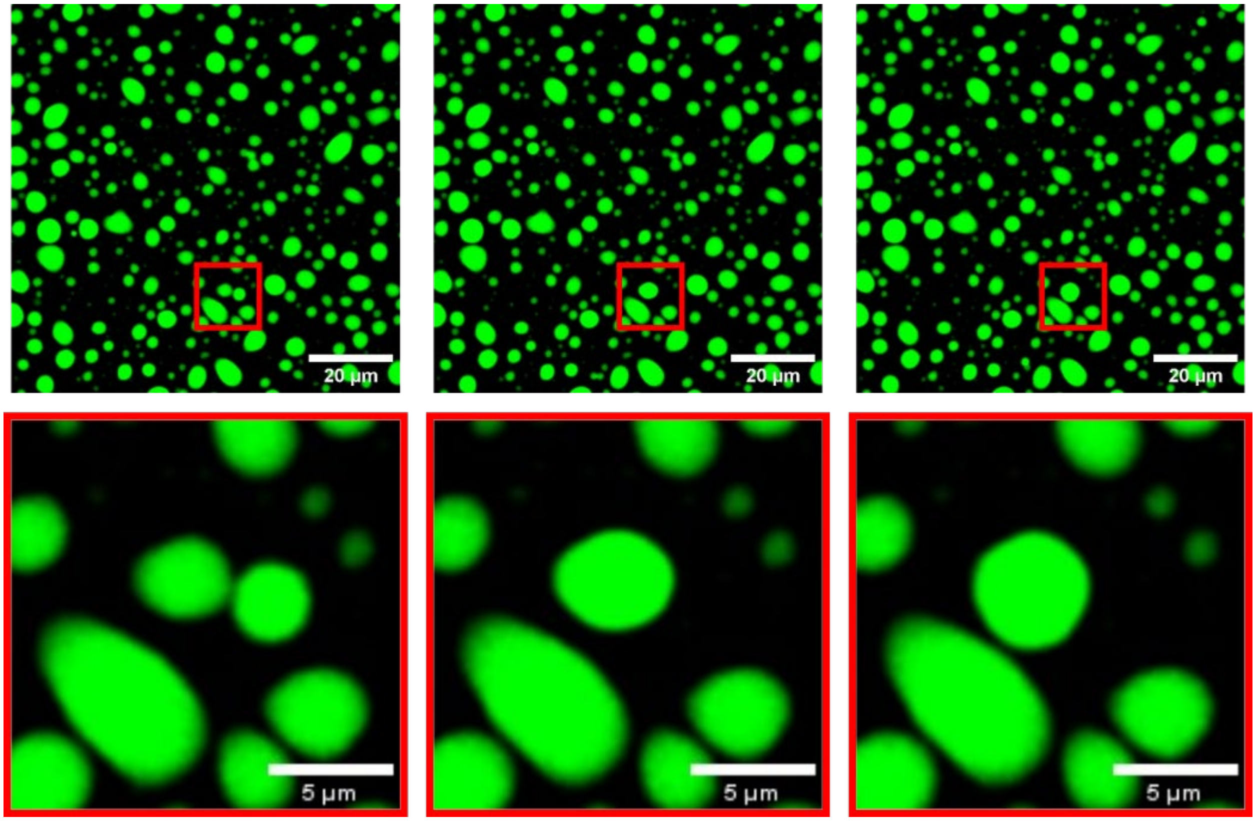


***Figure S5***. *CLSM images of 1% Alexa Fluor™ 488-labeled Nup (total concentration 0.165mM) in pure Nup98 condensates (NO PEG) captured 1 min after initializing LLPS. Images highlight the phenomenon of droplet merging. Bottom row are magnified views of the red squares in the top row.*

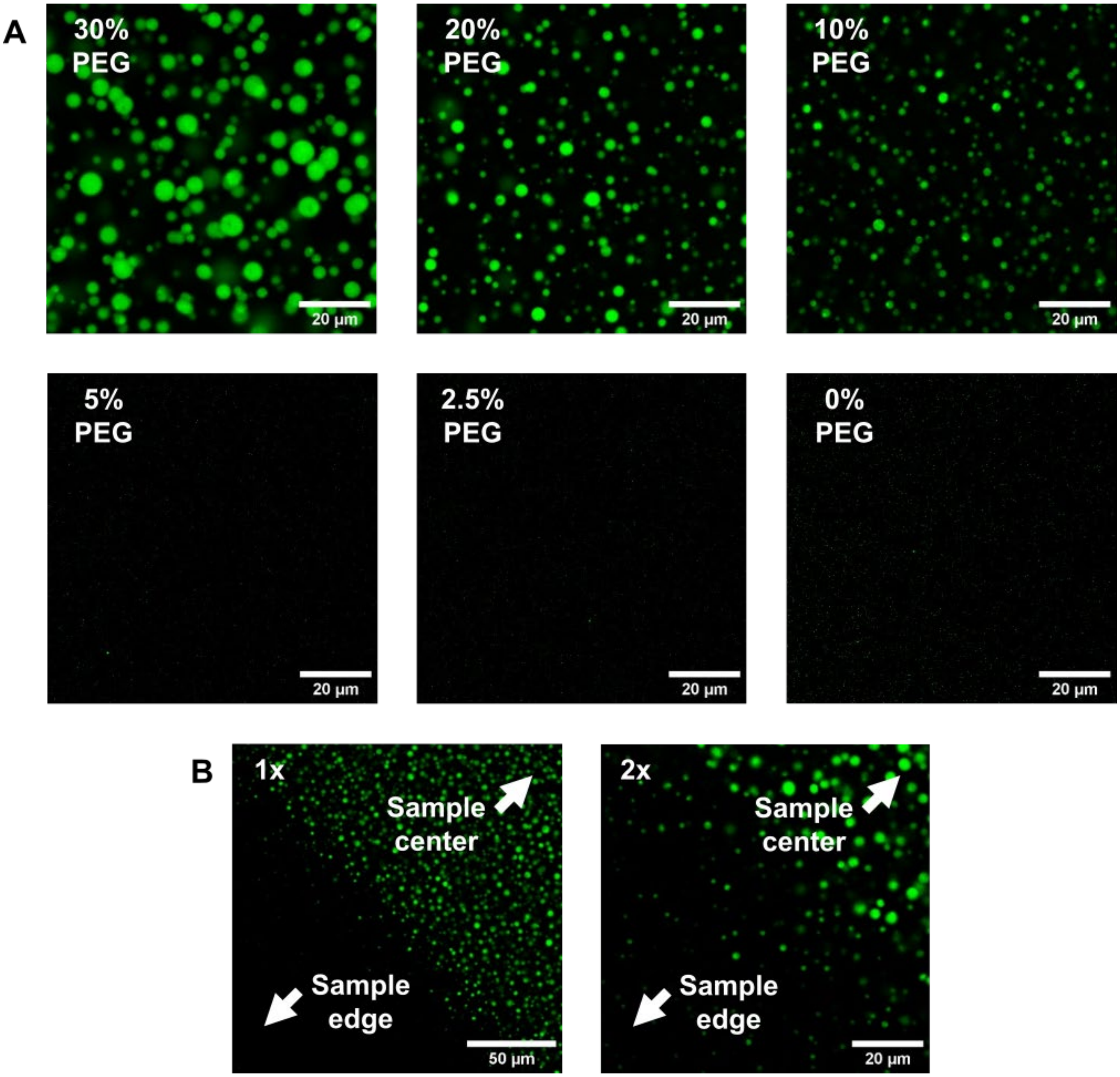


***Figure S6***. *CLSM images of 1% FITC-labeled BSA (total concentration 0.125mM) in BSA:PEG condensates.* ***A)*** *Images for 0%-30% PEG systems captured 5 min after initializing LLPS in the center of samples,* ***B)*** *Images for 30% PEG system captured 5 min after initializing LLPS focusing on a specific spot at the pre-existing interface of the BSA:PEG liquids, formed during the pipetting process.*

**SECTION S7**.

**The law of conservation of mass**.

According to mass conservation the following expressions should hold for the total volumes of components A (BSA) and B (PEG):

$$\bar{\phi}_A V = \phi_A^{(\alpha)} v^{(\alpha)} + \phi_A^{(\beta)} v^{(\beta)} \tag{S1}$$

$$\bar{\phi}_B V = \phi_B^{(\alpha)} v^{(\alpha)} + \phi_B^{(\beta)} v^{(\beta)} \tag{S2}$$

, with $\bar{\phi}_{i=A,B}$ the (known) mean volume fractions and $V$, $v^{(\alpha)}$ and $v^{(\beta)}$ the (unknown) total sample volume and respective phase volumes. Of the latter, $\alpha$ and $\beta$ respectively indicate the continuous and droplet phase. Furthermore, $\phi_{i=A,B}^{(j=\alpha,\beta)}$ denotes the volume fraction of a given component in a given phase. Of these, we assume $\phi_A^{(\alpha)} = 0$ and $\phi_A^{(\alpha)}$ and $\phi_B^{(\beta)}$ known (measured).

Dividing left and right by $V$ gives:

$$\bar{\phi}_A = \phi_A^{(\alpha)} \psi^{(\alpha)} + \phi_A^{(\beta)} \psi^{(\beta)} \tag{S3}$$

$$\bar{\phi}_B = \phi_B^{(\alpha)} \psi^{(\alpha)} + \phi_B^{(\beta)} \psi^{(\beta)} \tag{S4}$$

, with $\psi^{(j=\alpha,\beta)}$ the volume fractions of the phases. Since we have two-phase coexistence, we may write: $\psi^{(\alpha)} = 1 - \psi^{(\beta)}$. This, together with the assumption of zero protein content in the continuous gives two equations with two unknowns:

$$\bar{\phi}_A = \phi_A^{(\beta)} \psi^{(\beta)} \tag{S5}$$

$$\bar{\phi}_B = \phi_B^{(\alpha)} \left(1 - \psi^{(\beta)}\right) + \phi_B^{(\beta)} \psi^{(\beta)} \tag{S6}$$

Substitution of (S5) into (S6) eliminates $\psi^{(\beta)}$ and gives the following expression for the volume fraction of PEG in the continuous phase, as a function of the mean concentrations and the composition of the droplets:

$$\phi_B^{(\alpha)} = \frac{\bar{\phi}_B \phi_A^{(\beta)} - \bar{\phi}_A \phi_B^{(\beta)}}{\phi_A^{(\beta)} - \bar{\phi}_A} \tag{S7}$$

**Fig. S7** provides a summary of measured concentrations of BSA and PEG in the continuous phase and compares these findings with theoretical predictions based on the law of conservation of mass. The calculations presuppose that the protein is predominantly located in the droplet phase, with its minimal presence in the continuous phase deemed negligible. Theoretical values for the continuous phase were computed under the assumption that the BSA and PEG theoretical concentrations in the droplet phase aligns with experimentally derived values, thereby serving as a reference for mass distribution in the droplet phase. Columns highlighted in bold are intended for direct comparison.

| PEG content | Component | Introduced mass [mg] | Droplet phase concentration [mg·ml⁻¹] | **Continuous phase concentration [mg·ml⁻¹]** | Teoretical mass of components in droplet phase [mg] | Teoretical mass of components in continuous phase [mg] | Droplet phase theoretical concentration [mg·ml⁻¹] | **Expected continuous phase [mg·ml⁻¹]** |
|---|---|---|---|---|---|---|---|---|
| 10% PEG | PEG | 1000 | 51.72 | **91.26** | 7.96 | 992.04 | 51.72 | **91.47** |
| | BSA | 8.25 | 53.59 | **N.A.** | 8.25 | 0 | 53.59 | **0** |
| 20% PEG | PEG | 2000 | 88.17 | **167.92** | 8.66 | 1991.34 | 88.17 | **182.66** |
| | BSA | 8.25 | 83.95 | **N.A.** | 8.25 | 0 | 83.95 | **0** |
| 30% PEG | PEG | 3000 | 108.90 | **247.82** | 6.06 | 2993.94 | 108.90 | **273.56** |
| | BSA | 8.25 | 148.30 | **N.A.** | 8.25 | 0 | 148.30 | **0** |

***Figure S7**. Summary of BSA and PEG concentration and a comparison to theoretical values for the droplet and the continuous phase.*

**Description of the derivation of equations 8 and 9 included in the model section.**

We assume a solution of a sticky polymer. The stickers can form pairwise complexes, as well as trimers according to:

$$A + A \rightleftarrows A_2 \tag{S8}$$

$$A_2 + A \rightleftarrows A_3 \quad \text{(S9)}$$

We derive a partition function analogous to the derivation by Semenov and Rubinstein for pairwise associating stickers [S1] . The expression for the combinatorial factor for the possible sticker arrangements $P_{\text{comb}}$ is:

$$P_{\text{comb}} = \frac{n_{ST}!}{(n_{ST}-2n_P-3n_T)!n_P!n_T!2^{n_P}3^{2n_T}} \quad \text{(S10)}$$

, with $n_{ST}$, $n_P$ and $n_T$ the total number of stickers, dimers and trimers, respectively. The probability that all $2n_P$ and $3n_T$ stickers are in each other vicinity to for the respective complexes, is given by:

$$W = \left(\frac{v_P}{V}\right)^{n_P} \left(\frac{v_T}{V}\right)^{2n_T} \quad \text{(S11)}$$

, with $v_P$ and $v_T$ the volume of a binary and a ternary complex. The sticker partition function due to sticker association is given by:

$$Z_{ST} = P_{\text{comb}} W \exp(n_P \varepsilon_1) \exp\big(n_T(\varepsilon_1 + \varepsilon_2)\big) \quad \text{(S12)}$$

, with $\varepsilon_1$ and $\varepsilon_2$ the energy in units of $k_B T$ associated with the stepwise formation a binary and a ternary complex. The sticker contribution to the free energy density is:

$$\frac{F_{ST}}{k_B T V} = \frac{f_{ST}}{k_B T} = -\frac{1}{V}\ln Z_{ST} = -\frac{1}{V}\ln(P_{\text{comb}} W) - \frac{1}{V}\big(n_P \varepsilon_1 + n_T(\varepsilon_1 + \varepsilon_2)\big) \quad \text{(S13)}$$

, with $F_{ST}$ the sticker free energy and $V$ the total volume of the system. Substituting **Eq. S10 and S12** and applying Stirling's approximation yields:

$$\frac{f_{ST}}{k_B T} = -\frac{c}{l}\left[\frac{p_1}{2}\ln\left(\frac{c v_P}{el}\right) + \frac{2p_2}{3}\ln\left(\frac{c v_T}{el}\right)\right] + \frac{c}{l}\left[\frac{p_1}{2}\ln(p_1) + \frac{p_2}{3}\ln(3p_2) + (1-p_1-p_2)\ln(1-p_1-p_2)\right] - \frac{p_1 c}{2l}\varepsilon_1 - \frac{p_2 c}{3l}(\varepsilon_1 + \varepsilon_2) \quad \text{(S14)}$$

, with: $p_1 = 2n_P/n_{ST}$, $p_2 = 3n_T/n_{ST}$, $n_{ST} = \frac{c}{l}V$, $c$ the monomer number density and $l$ the number of monomers between two stickers. Upon substituting: $c = \Phi/v_0$ with $\Phi$ the volume fraction of the stickerbearing polymer and $v_0$ the volume of a monomer or grid site, as well as the relation: $cv_{i=P,T} = cx_i v_0 = \Phi x_i$ with $x_i$ a dimensionless multiplication factor, we obtain:

$$\frac{f_{ST}}{k_B T} = -\frac{\Phi}{l v_0}\left[\frac{p_1}{2}\ln\left(\frac{\Phi x_P}{el}\right) + \frac{2p_2}{3}\ln\left(\frac{\Phi x_T}{el}\right)\right] + \frac{\Phi}{l v_0}\left[\frac{p_1}{2}\ln(p_1) + \frac{p_2}{3}\ln(3p_2) + (1 - p_1 - p_2)\ln(1 - p_1 - p_2)\right] - \frac{p_1\Phi}{2l v_0}\varepsilon_1 - \frac{p_2\Phi}{3l v_0}(\varepsilon_1 + \varepsilon_2) \quad \text{(S15)}$$

Realizing that the total volume is given by the product of the total number of lattice sites, multiplied by $v_0$, we can eliminate the latter and write the dimensionless free energy on a per lattice site basis as:

$$f_{\text{bind}} = -\frac{\Phi}{l}\left[\frac{p_1}{2}\ln\left(\frac{\Phi x_P}{el}\right) + \frac{2p_2}{3}\ln\left(\frac{\Phi x_T}{el}\right)\right] + \frac{\Phi}{l}\left[\frac{p_1}{2}\ln(p_1) + \frac{p_2}{3}\ln(3p_2) + (1 - p_1 - p_2)\ln(1 - p_1 - p_2)\right] - \frac{p_1\Phi}{2l}\varepsilon_1 - \frac{p_2\Phi}{3l}(\varepsilon_1 + \varepsilon_2) \quad \text{(S16)}$$

Assuming the bound sticker fraction to be in equilibrium (see main text), subtracting the free energy of the pure states and assuming $x_P/x_T = 2/3$ yields the set of **Eq. 8, 9, 10** after some algebra. We finally note that in our model the association constants $K_1$ and $K_2$ for equilibria (**Eq. S8 and S9**), given in **Eq. 9 and 10**, are related to the binding energies $\varepsilon_1$ and $\varepsilon_2$ according to:

$$K_1 = \frac{1}{2} x_P v_m \exp(\varepsilon_1) \quad \text{(S17)}$$

$$K_2 = \frac{1}{3} x_T v_m \exp(\varepsilon_2) \quad \text{(S18)}$$

Here, with $v_m = \mathcal{N}_A v_0$ a reference volume, being the monomer molar volume, with $\mathcal{N}_A$ Avogadro's number. For all calculations we set $v_m$ = 1 M$^{-1}$.

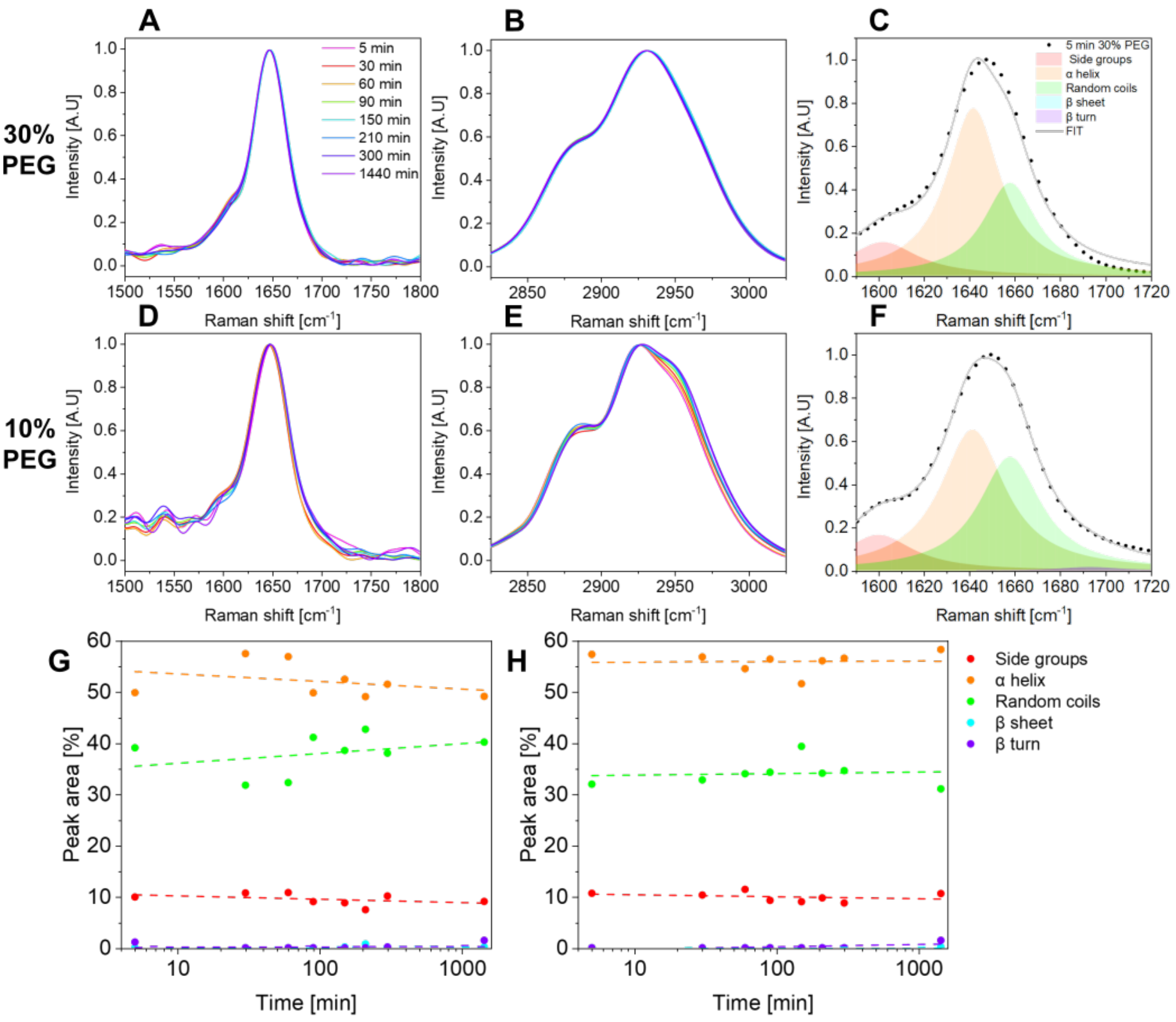


***Figure S8***. *Normalized BCARS fingerprint (**A** and **D**) and $CH_x$ (**B** and **E**) spectra comparison for droplet phase of BSA:PEG system at various PEG concentrations over progressive maturation time. Amide I spectra deconvolution for **C**) 30% PEG system at 5min and **F**) 10% PEG system at 5min. Contribution of deconvoluted peaks area over time for **G**) 10% PEG and **H**) 30% PEG systems.*

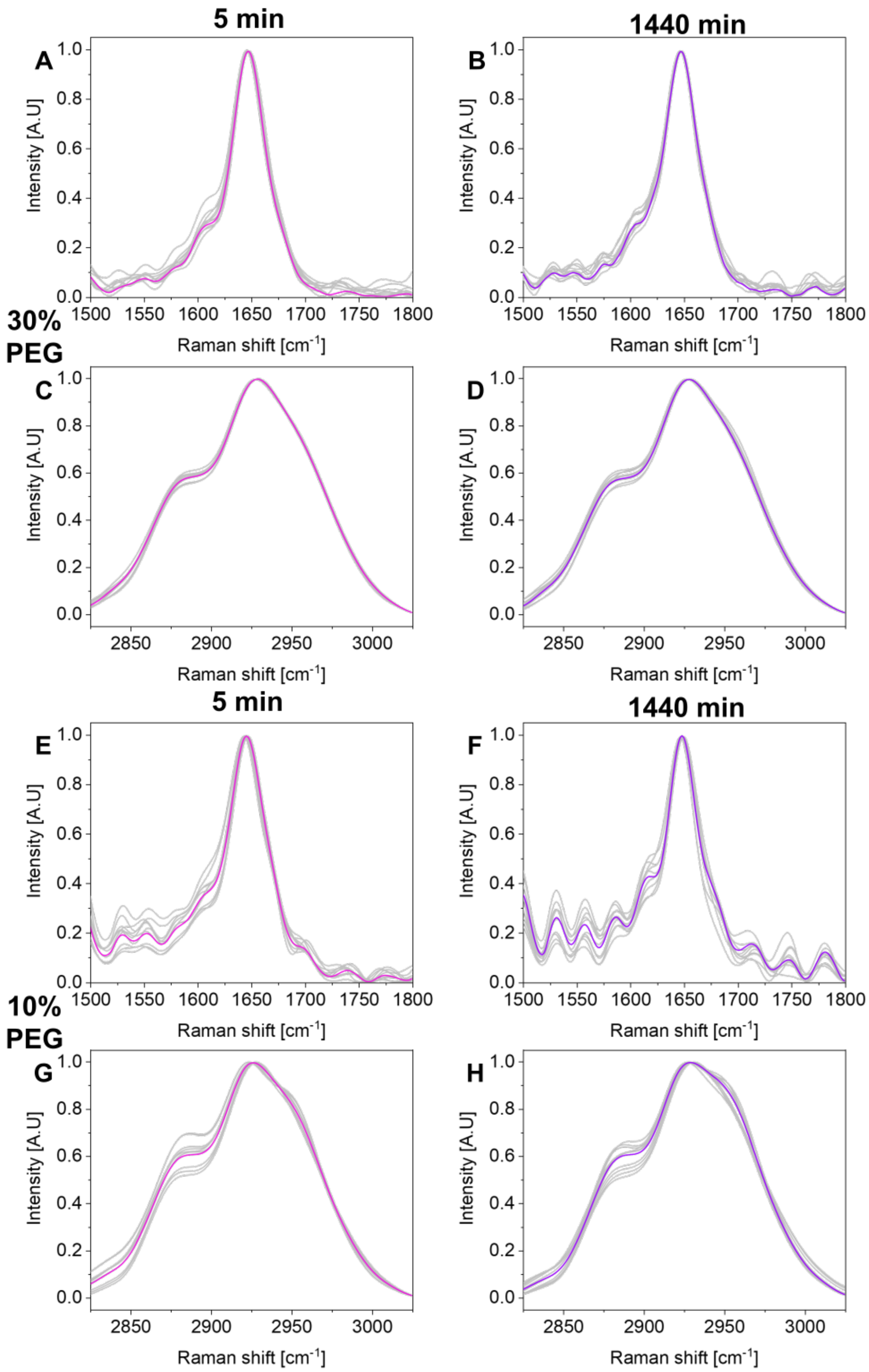


***Figure S9***. *Normalized BCARS fingerprint (**A**, **B**, **E**, **F**) and $CH_x$ (**C**, **D**, **G**, **H**) spectra comparison for droplet phase of BSA:PEG system for 30% PEG (A, B, C, D) and 10% PEG (E, F, G, H) at 5 min (A, C, E, G) and 1440 min (B, D, F, H). Color lines are averaged data. Grey curves represent 10 exemplary data curves each integrated from >5 individual pixels.*

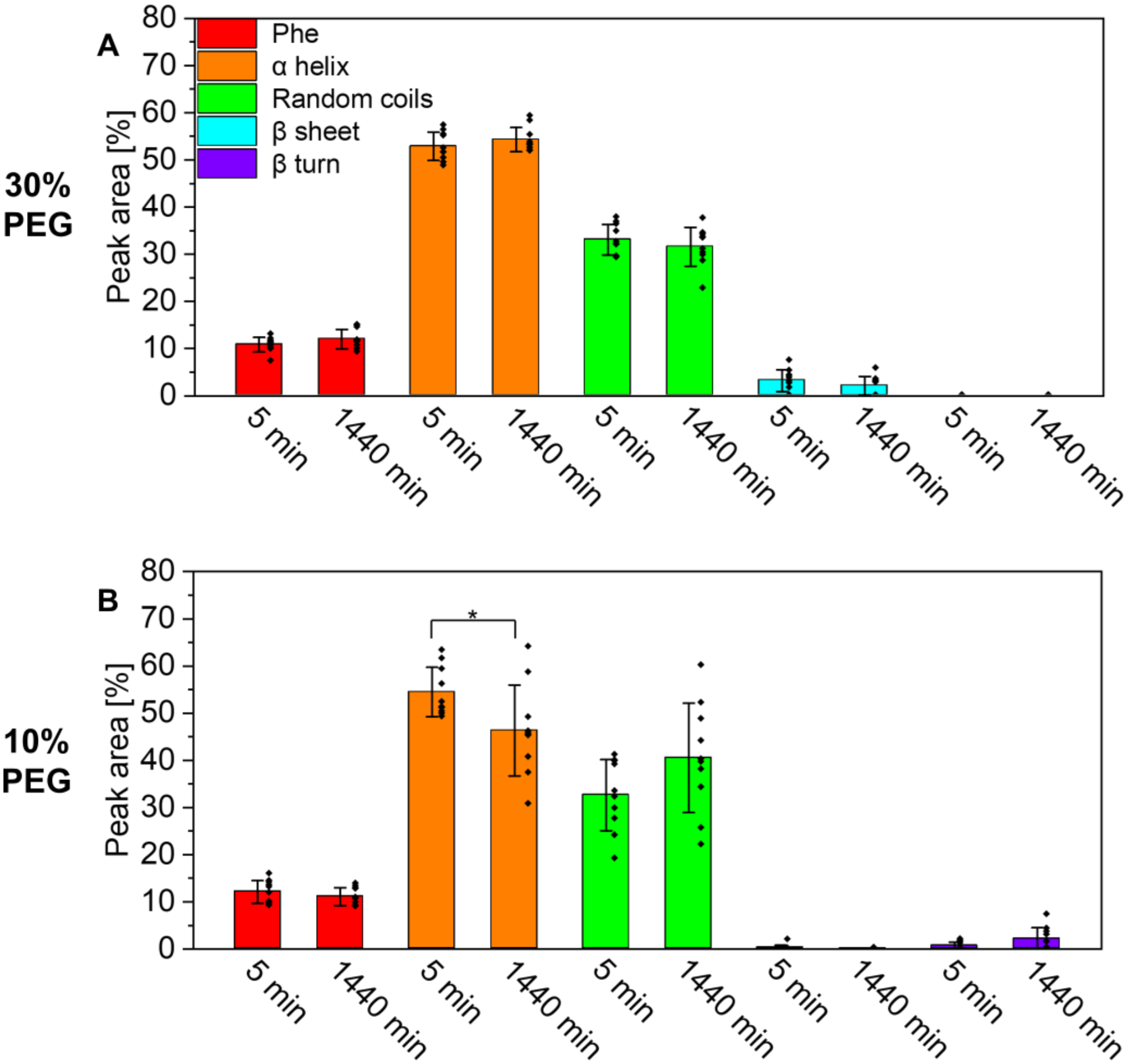


***Figure S10**. Contribution of subpeaks in Amide I deconvoluted spectra of BSA:PEG systems **A**) 30% PEG and **B**) 10% PEG at 5 min and 1440 min. Sample population matches S9; n= 10; error bars are std; asterisks symbolize statistical difference of mean values (one-way ANOVA p<0.05); black dots are unique data points.*

[S1] A. N. Semenov and M. Rubinstein, "Thermoreversible gelation in solutions of associative polymers. 1. statics," *Macromolecules*, vol. 31, no. 4, pp. 1373–1385, 1998. doi:10.1021/ma970616h
